# Comparative analysis of active sites in P-loop nucleoside triphosphatases suggests an ancestral activation mechanism

**DOI:** 10.1101/439992

**Authors:** Daria N. Shalaeva, Dmitry A. Cherepanov, Michael Y. Galperin, Armen Y. Mulkidjanian

**Author notes:** **For correspondence:** Armen Y. Mulkidjanian, School of Physics, University of Osnabrück, 16 D-49069, Osnabrück, Germany. E-mail addresses: Daria N. Shalaeva, Dmitry A. Cherepanov, Michael Y. Galperin, Armen Y. Mulkidjanian.

## Abstract

P-loop nucleoside triphosphatases (NTPases) share common Walker A (P-loop) and Walker B sequence motifs and depend on activating moieties (Arg or Lys fingers or a K^+^ ion). In search for a common catalytic mechanism, we combined structure comparisons of active sites in major classes of P-loop NTPases with molecular dynamics (MD) simulations of the Ras GTPase, a well-studied oncoprotein. Comparative structure analysis showed that positively charged activating moieties interact with gamma-phosphate groups of NTP substrates in all major classes of P-loop NTPases. In MD simulations, interaction of the activating Arg finger with the Mg-GTP-Ras complex led to the rotation of the gamma-phosphate group by 40 degrees enabling its interaction with the backbone amide group of Gly13. In all analyzed structures, the residue that corresponds to Gly13 of Ras was in a position to stabilize gamma-phosphate after its rotation, suggesting a common ancestral activation mechanism within the entire superfamily.

## Introduction

Hydrolysis of nucleoside triphosphates (NTPs), such as ATP or GTP, by various nucleoside triphosphatases (NTPases) is a key reaction in the cell. Most widespread are the P-loop NTPases that make up 10-20% gene products in a typical cell (Koonin et al., 2000; Leipe et al., 2002). P-loop domains appear to be some of the most ancient protein domains dating back to the Last Universal Cellular Ancestor (LUCA) (Alva et al., 2015; Lupas et al., 2001; Orengo & Thornton, 2005; Ponting & Russell, 2002; Ranea et al., 2006; Söding & Lupas, 2003). They are found in proteins that hydrolyze ATP to perform mechanical work, such as kinesin, myosin and dynein; rotary ATP synthases; DNA and RNA helicases, many GTPases, including α-subunits of signaling G-proteins, and other enzymes. While the expansion of most protein families correlates with expansion of their catalytic repertoire, P-loop NTPases are unusual in being abundant but catalyzing only a few types of hydrolytic reactions (Anantharaman et al., 2003).

The P-loop fold is a variation of the Rossman fold, a 3-layer αβα sandwich (Leipe, et al., 2002). One of the loops contains the GxxxxGK[ST] sequence motif, known as the Walker A motif (Walker et al., 1982). This motif is responsible for binding the NTP’s triphosphate chain and is often referred to as the P-loop (<u>*p*</u>hosphate-binding loop) motif (Saraste et al., 1990). The conserved Lys residue of the P-loop forms hydrogen bonds (H-bonds) with β- and γ-phosphate groups of the NTP. The Ser/Thr residue coordinates the Mg^2+^ ion, which also binds between β- and γ-phosphates, but from the other side (Fig. 1A). The Walker B motif *hhhh*D, where ‘*h’* denotes a hydrophobic residue, provides the conserved Asp residue that usually serves as an additional Mg^2+^ ligand (Walker, et al., 1982).

**Figure 1.**
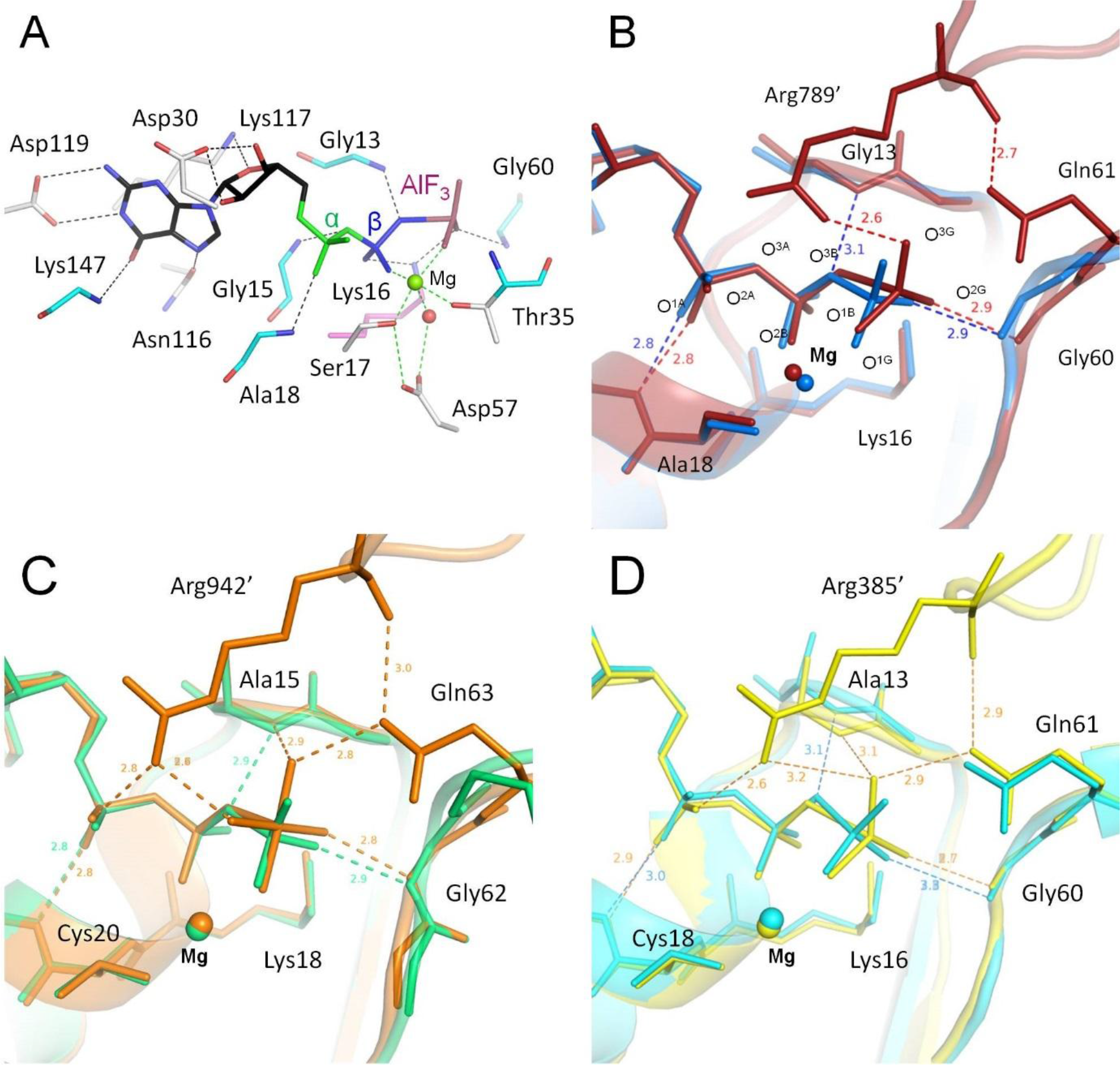
Phosphate chain conformations in Mg-NTP complexes. **A.** Structure of the transition state analog GDP-AlF_3_ bound to the complex of H-Ras GTPase [PDB: 1WQ1]. The topology of the bonds was taken from (Wittinghofer & Vetter, 2011). Nucleotides, their analogs, and adjacent amino acid residues are shown as sticks, P-loop lysine (Lys16) is shown in pink, backbone groups are in cyan, carbon atoms of GDP are in black, phosphate groups and their analogs are colored as follows: α-phosphate in green, β-phosphate in blue, and γ-phosphate analog AlF_3_ in dark red. Mg^2+^ ion is shown as a green sphere, the catalytic water molecule as a red sphere. Black dashed lines indicate hydrogen bonds and salt bridges, green dashed lines indicate interactions of Mg^2+^ ion coordination sphere. **B.** Superposition of crystal structures of Ras and Ras/RasGAP complex. The structure of the Ras with bound GTP [PDB: 1QRA] is shown in blue, the structure of the Ras/RasGAP complex with the transition state analog GDP-AlF_3_ [PDB: 1WQ1] is shown in red. The oxygen atoms are named according to the CHARMM naming scheme (Vanommeslaeghe, et al., 2010) and the IUPAC recommendations (Blackburn, et al., 2017). Protein backbones are shown as cartoons; GTP, its analogs and surrounding amino acid residues are shown as sticks, Mg^2+^ ions are shown as spheres. Dashed lines indicate hydrogen bonds or coordination bonds shorter than 3.4 Å, all distances are in Å. **C.** Superposition of crystal structures of RhoA and RhoA/RhoGAP complex. The structure of RhoA with bound GTP [PDB: 4XOI] is shown in green, the structure of the RhoA/RhoGAP complex with the transition state analog GDP-AlF_4_ [PDB: 5JCP] is in orange. Other designations as in Fig. 1B. **D.** Superposition of crystal structures of Ras-like GTPase Cdc42 and Cdc42/MgcRacGAP complex. The structure of the Cdc42 with bound GMP-PCP (non-hydrolyzable analog of GTP) [PDB: 2QRZ] is shown in cyan, the structure of the Cdc42/MgcRacGAP complex with the transition state analog GDP-AlF_3_ [PDB: 5C2J] is in yellow. Other designations as in Fig. 1B.

The best-studied examples of P-loop NTPases are small regulatory GTPases that include H-, N-and K-Ras (<u>ra</u>t <u>s</u>arcoma) proteins. Human Ras GTPases get converted into active oncoproteins by mutations in positions 12, 13, or 61 (Bos et al., 1987; Prior et al., 2012; Wey et al., 2013). NTP hydrolysis is achieved through interaction of the P-loop domain with another protein or domain, which provides an activating moiety, usually a Lys or Arg “finger”, that gets inserted into the active site and promotes catalysis (Ash et al., 2012; Jin, Molt, et al., 2017; Kamerlin et al., 2013; Ogura et al., 2004; Scheffzek et al., 1997; Scrima & Wittinghofer, 2006; Wendler et al., 2012; Wittinghofer & Vetter, 2011). In Ras and related GTPases, the Arg finger, which triggers the hydrolysis and cancels the oncogenic signal, comes from specific <u>G</u>TPase <u>a</u>ctivating <u>p</u>roteins (GAPs) (Scheffzek, et al., 1997; Wittinghofer & Vetter, 2011).

Cleavage of the NTP phosphoanhydride bond results from the nucleophilic attack by a polarized water molecule that is believed to be coordinated by so-called “catalytic/essential” residues, see (Jin, Molt, et al., 2017; Jin, Richards, et al., 2017; Kamerlin, et al., 2013; Kiani & Fischer, 2016) for reviews. The exact catalytic functions of such residues mostly remain unclear.

It also remains unclear whether there is some universal catalytic mechanism common for all P-loop NTPases, see (Kamerlin, et al., 2013; Ogura, et al., 2004; Wittinghofer, 2006). Gerwert and colleagues proposed that, during GTP hydrolysis by Ras and heterotrimeric G proteins, the Arg finger rotates the α-phosphate towards an eclipsed conformation with respect to β- and γ-phosphates, which would favor the bond cleavage (Gerwert et al., 2017; Mann et al., 2016; Rudack et al., 2012). Kiani and Fischer emphasized the possible participation of Glu residues in polarizing the attacking water molecule in myosin, kinesin and rotary ATPase (Kiani & Fischer, 2016). Blackburn and colleagues proposed that the insertion of the activating Arg residue leads to the reshuffling of the hydrogen bonded network, which drives the displacement of the attacking water molecule into the reactive position (Jin, Molt, et al., 2017; Jin et al., 2016; Jin, Richards, et al., 2017).

Since in some P-loop NTPases, the role of activating Arg/Lys residues is played by K^+^ ions (Anand et al., 2010; Ash, et al., 2012; Bohme et al., 2010; Meyer et al., 2009; Scrima & Wittinghofer, 2006), we have combined, in a separate study, structural analysis of diverse P-loop NTPases with molecular dynamics (MD) simulations of the Mg^2+^-ATP complex in water in the presence of monovalent cations (Shalaeva et al., submitted). Monovalent cations were found to occupy two specific positions around the Mg^2+^-ATP complex. Namely, cations bound between β- and γ-phosphates (the BG site), in the position of the conserved Lys residue of the P-loop (*cf* Fig. 1A), and between α- and γ-phosphates (the AG site), in the position of the activating Arg/Lys residue. Additionally, we observed weak cation binding in the proximity of γ-phosphate (the G-site), which matches the location of the catalytic/essential residues in P-loop NTPases. The catalytically prone conformation of the phosphate chain in the active sites of diverse P-loop NTPases was found to be similar to that observed in water in the presence of K^+^ and NH_4_^+^ ions and more stretched than the one observed in the presence of the smaller Na^+^ ions (Shalaeva et al., submitted). We concluded that the shape of the phosphate chain in the active sites of P-loop NTPases and the catalytically favorable electrostatic environment are defined by certain inherent properties of Mg^2+^-NTP complexes in the presence of large monovalent cations. The conservation of the electrostatic environment of the NTP substrate in different classes of P-loop ATPases and GTPases could underlie the long-sought basal mechanism of hydrolysis common to all P-loop ATPases and GTPases.

Here, we used evolutionary tools to look for mechanistic traits shared by most P-loop NTPases. By combining comparative structural analysis of major families of P-loop NTPases with MD simulations of human Ras GTPase, we identified several structural traits that are shared by catalytic sites in diverse families of P-loop NTPases and appear to be inherited from the ancestral form(s) of P-loop proteins. These traits imply a common catalytic mechanism in P-loop NTPases. The proposed mechanism assigns a key role to the rotation of γ-phosphate and suggests that the backbone nitrogen atom of a specific residue of the P-loop (Gly13 in Ras, whose mutations often lead to cancer) provides an H-bond to stabilize γ-phosphate after its rotation. In addition, from our MD simulation data, we estimated the free energy needed for inserting the activating moiety and twisting the gamma-phosphate group as 20-25 kJ/mol. This energy appears to be derived from the interaction between the P-loop domain and the activating protein domain. This way, NTP hydrolysis can be controlled through specific protein-protein interactions, which explains the abundance and variety of P-loop NTPases despite their seemingly narrow catalytic repertoire.

## Results

### In the Ras/RasGAP complex, the Arg finger rotates the γ-phosphate group

The structures of P-loop NTPases with bound ATP or GTP are virtually absent in the Protein Data Bank (PDB) because of the rapid hydrolysis of NTPs. Therefore, most studies utilize structures with either non-hydrolyzable analogs of NTPs or transition state analogs, such as ADP/GDP complexes with BeF_3_^−^, AlF_3_, AlF_4_^−^, MgF_3_^−^ see Fig. 1A, (Jin, Molt, et al., 2017; Jin, Richards, et al., 2017; Scheffzek, et al., 1997) and (Shalaeva et al., submitted). Human Ras GTPase (P21/H-RAS-1) and some related proteins are rare exceptions, whose structures with bound Mg-GTP could be obtained due to their low intrinsic activity in the absence of the respective activating proteins (Kotting & Gerwert, 2004; Shutes & Der, 2006). For Ras GTPase, only one such structure [PDB: 1QRA] (Scheidig et al., 1999) is a structure of the wild-type protein, all others include mutations of residues in the proximity of the substrate binding site (Table S1). For the Ras/RasGAP complex, a wild-type structure is also available, in complex with a transition state analog GDP-AlF_3_ [PDB: 1WQ1] (Scheffzek, et al., 1997). Superposition of these two structures showed that the AlF_3_ group, which mimics the γ-phosphate, is rotated and shifted as compared to the position of γ-phosphate in the GTP, while the rest of the substrate molecules aligns almost perfectly (Fig. 1B and Table S2). This rotation is accompanied by a displacement of the Mg atom, which indicates certain elasticity of the bonding network around the γ-phosphate.

The certain flexibility of γ-phosphate follows also from superposition of numerous structures of Ras proteins with GTP and its analogs (Fig. S1A, S1B). Extensive comparative structure analysis of the transition state-like complexes of diverse P-loop NTPases, conducted by Blackburn and coworkers, also showed highly variable orientations of the γ-phosphate mimicking groups (Jin, Molt, et al., 2017; Jin, Richards, et al., 2017), indicating the possibility of γ-phosphate rotation.

Another structure with a bound GTP molecule is available for the Ras-like GTPase RhoA [PDB: 4XOI] (Sun et al., 2015). In Fig. 1C, this structure is superposed with the structure of RhoA ([PDB: 5JCP] (Bao et al., 2016)) in complex with its activating protein RhoGAP and GDP-AlF_4_^−^. The planar AlF_4_^−^ ion is believed to be the closest analog of the transition state (Jin, Richards, et al., 2017). Superposition of the two RhoA structures highlighted not only the conformational flexibility of the terminal group, but also the changes in the hydrogen bond network around it, particularly the formation of a new H-bond between the backbone nitrogen of Ala15 (which corresponds to Gly13 of Ras) and one of the F atoms of AlF_4_^−^. Being intrigued by this H-bond, we have checked for its presence in structures of P-loop NTPases that contain ADP or GDP in complex with AlF_4_^−^. Using InterPro protein domain annotations and RCSB PDB search engine, we have identified 60 X-ray structures of P-loop proteins in complex with GDP-AlF_4_^−^ or ADP-AlF_4_^−^, which include in total 130 individual nucleotide-protein complexes (Table S3). In 105 of such complexes, the backbone nitrogen of the Gly13 (or another residue in that position) is within 3.3 Å of the nearest fluorine atom, indicating a possible H-bond (Howard et al., 1996). The exceptions were myosines, the adenylate kinase and replicative DNA helicase, for which the distances were in the range of 3.5–4.6 Å.

Encouraged by finding numerous bonds between the AlF_4_^−^ moieties and counterparts of Gly13 of Ras, we revisited the AlF_3_-containing structures of P-loop NTPases. Using the same protocol as with AlF_4_-containing structures, we found 34 PDB entries of P-loop proteins containing ADP-AlF_3_ or GDP-AlF_3_ complexes, with a total of 58 individual protein-nucleotide complexes, see Table S4. Among them, four PDB entries contain fluoride atom of AlF_3_ within 3.3 Å of the backbone nitrogen of a residue that corresponds to Gly13 of Ras. Those structures are Cell division control protein 42 homolog (PDB: 1GRN), nitrogenase-like ATPase ChlL (PDB: 2YNM), flagellar biosynthesis ATPase FlhF (PDB: 3SYN) and thymidylate kinase (PDB: 1E2E). Formation of an H-bond between the counterpart of Gly13 of Ras and the oxygen of γ-phosphate is also seen in the structure of the guanosine-5’-[(β,γ)-methyleno]triphosphate (GMP-PCP) containing Ras-like GTPase Cdc42 in complex with its activating Male Germ Cell RacGAP domain (Toure et al., 1998), but not in the structure of GMP-PCP-containing Cdc42 GTPase alone (Phillips et al., 2008), see Fig. 1D.

While structure comparisons operate with individual averaged snapshots of protein conformations, MD simulations yield ensembles of allowed protein conformations and enable estimation of the stability of particular interactions, including the H-bonds. The possibility of γ-phosphate rotation and its interaction with counterparts of Gly13 of Ras, evident from the structure comparison (Table S3, S4, Fig. 1B-D, S1A, S1B), prompted us to perform MD simulations of the Ras and Ras/RasGAP complex to investigate further the substrate-protein interactions, as well as the mobility of the γ-phosphate group.

In the GTP-containing structure of wild-type Ras ([PDB: 1QRA], Fig. 1B), the so-called Switch I region of the catalytic pocket is in the “open” state (Fig. S1B) with Tyr32 turned away from the NTP-binding site, whereas the “closed” state, where Tyr32 makes a H-bond with O^3G^ oxygen atom, is considered more probable for Ras in the absence of RasGAP (Wittinghofer & Vetter, 2011). Thus, for MD simulations, we used the high-resolution structure of wild-type Ras with non-hydrolyzable analog GMP-PNP bound in the closed state [PDB: 5WDO] (Scheidig, et al., 1999). As a starting structure for the H-Ras/RasGAP complex, we used the same structure as in Fig. 1B, namely the crystal structure [PDB: 1WQ1] of the Ras/RasGAP complex with the transition state analog GDP-AlF_3_ (Scheffzek, et al., 1997). In both structures, we replaced the nucleotide analogs with a GTP molecule using the coordinates taken from the wild-type GTP-Ras complex [PDB: 1QRA], after superposition of the P-loop regions of the Ras protein. The resulting structures of the Mg-GTP-Ras and Mg-GTP-Ras/RasGAP complex were then used for MD simulations, as described in Methods. Representative structures of both Mg-GTP-Ras and Mg-GTP-Ras/RasGAP complexes, as sampled from free MD simulations after equilibration of the systems, are available on request.

MD simulations showed that the bonds that stabilize the phosphate chain of the bound Mg-GTP in H-Ras (Fig. 1A) persisted upon formation of the Ras/RasGAP complex (Table 1, Fig. 2, S3). These interactions involved each of the three phosphate groups: while O^1A^ formed an H-bond with the backbone nitrogen of Ala18, O^1B^ and O^2G^ atoms were coordinated by the amino group of Lys16 (Fig. 2A, hereafter, the atom names follow the CHARMM naming scheme (Vanommeslaeghe et al., 2010) and the recent IUPAC Recommendations (Blackburn et al., 2017), see Fig. 1B). The O^2B^ and O^1G^ atoms coordinated the Mg^2+^ ion, together with oxygen atoms of Ser17 of the P-loop motif, Thr35, and two water molecules. The oxygen atoms of the phosphate chain additionally formed several H-bonds with the backbone nitrogens of the P-loop residues: O^1B^ with Val14 and Gly15, and O^3B^ with Gly13 (Table 1, Fig. 1A, 2, S3).

**Table 1.** Interatomic distances and hydrogen bonding times in the GTP-binding sites of Mg-GTP-Ras and Mg-GTP-Ras/RasGAP complexes during MD simulations^a^

| Interacting atoms<sup>b</sup> |  | Mg-GTP-Ras |  | Mg-GTP-Ras-RasGAP |  |
| --- | --- | --- | --- | --- | --- |
| Donor | Acceptor | Average distance, Å | H-bonding time, % | Average distance, Å | H-bonding time, % |
| Gly13 N | O<sup>3B</sup> | 3.02±0.15 | 76.5 | 3.24±0.22 | 26.8 |
| Gly13 N | O<sup>2G</sup> | 9.91±0.25 | 0.2 | 3.29±0.37 | 39.3 |
| Val14 N | O<sup>1B</sup> | 3.11±0.16 | 16 | 3.09±0.16 | 15 |
| Gly15 N | O<sup>1B</sup> | 3.0±0.14 | 60.5 | 2.99±0.14 | 57.7 |
| Lys16 N | O<sup>1B</sup> | 2.99±0.14 | 84.5 | 2.94±0.13 | 92 |
| Lys16 NZ | O<sup>1B</sup> | 2.73±0.10 | 100 | 2.69±0.09 | 99.8 |
| Lys16 NZ | O<sup>2G</sup> | 2.57±0.06 | 100 | 2.58±0.06 | 100 |
| Ser17 N | O<sup>2B</sup> | 3.07±0.12 | 63 | 2.95±0.12 | 90.9 |
| Ala18 N | O<sup>1A</sup> | 2.8±0.10 | 99.5 | 2.92±0.14 | 90.5 |
| Tyr32 OH | O<sup>3G</sup> | 2.8±0.70 | 88.1 | 6.51±0.29 | 0 |
| Thr35 N | W<sub>cat</sub><sup>c</sup> | N/A<sup>c</sup> | 51.5 | 2.94±0.19 | 86.5 |
| Gly60 N | O<sup>2G</sup> | 2.77±0.12 | 97.3 | 2.92±0.23 | 73.6 |
| Gln61 NE | O<sup>3G</sup> | 8.81±1.49 | 0 | 3.05±0.27 | 69.6 |
| W<sub>cat</sub><sup>c</sup> | Gln61 OE | N/A<sup>c</sup> | 0.3 | 2.8±0.19 | 91.1 |
| Gln61 NE | Arg789<sup>d</sup> O | N/A<sup>d</sup> |  | 3.48±0.38 | 19.3 |
| Arg789<sup>d</sup> NH2 | Thr785<sup>d</sup> O |  |  | 2.82±0.13 | 95.4 |
| Arg789<sup>d</sup> NH1 | O<sup>2A</sup> |  |  | 2.83±0.19 | 95.7 |
| Arg789<sup>d</sup> NH1 | O<sup>3G</sup> |  |  | 2.73±0.10 | 99.7 |

<sup>a</sup> – Interactions between the atoms were evaluated from the conformations recorded from MD simulations at 25-ps intervals. Hydrogen bonding times were calculated as a fraction of conformations, in which (i) the distance between the hydrogen atom and the acceptor was less than 2.2 Å, and (ii) the angle between the donor, the hydrogen atom, and the acceptor was wider than 120° (Khrenova et al., 2014). Interatomic distances and angles (means and standard deviations) were calculated in MatLab.
<sup>b</sup> – Atom designations follow the PDB rules for protein residues and the CHARMM naming scheme (Vanommeslaeghe, et al., 2010) as well as the IUPAC recommendations (Blackburn, et al., 2017).
<sup>c</sup> – W<sub>cat</sub> indicates the water molecule in the catalytic site. In Mg-GTP-Ras simulations, these water molecules were drifting freely, rendering distance measurements meaningless.
<sup>d</sup> – Arg789 comes from RasGAP and was absent from the Mg-GTP-Ras simulations

**Figure 2.**
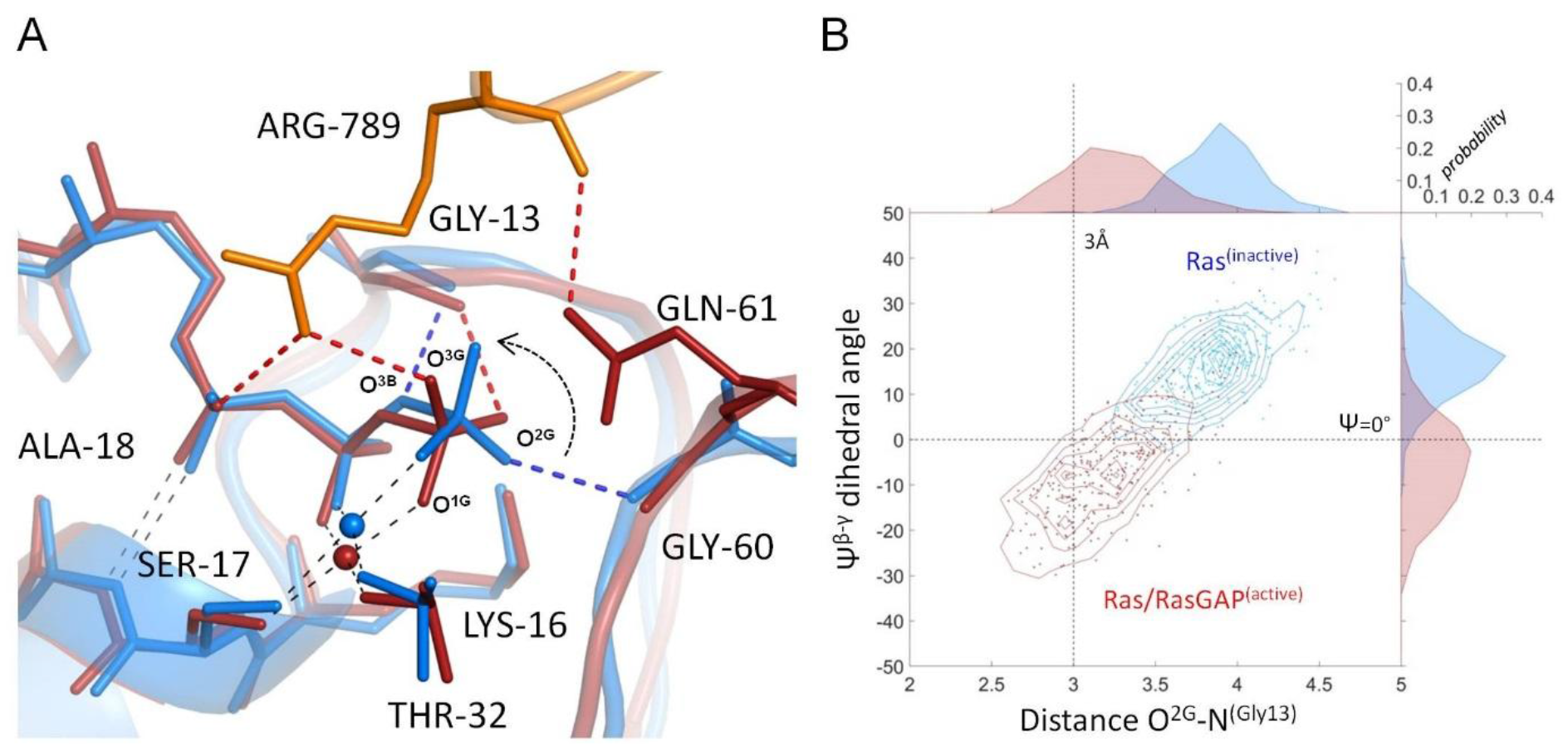
Molecular dynamics of GTP in the catalytic site of Ras GTPase. **A.** Superposition of Mg-GTP bound to Ras (all atoms shown in blue) with Mg-GTP bound to Ras/RasGAP (all Mg-GTP and Ras atoms shown in red, the RasGAP fragment is in orange); representative structures were sampled from a series of MD simulations. Black dashed lines indicate hydrogen bonds or coordination bonds that are present in both structures, red and blue dashed lines indicate H-bonds differing between the two structures. **B.** Conformational space of GTP in Ras (blue) and Ras/RasGAP (red) sampled from MD simulations and plotted as a scatter plot of the Ψ^β-γ^ dihedral angle(Y-axis) against the length of the hydrogen bond between O^2G^ and the backbone nitrogen of Gly13 (X-axis). Individual distribution histograms for the Ψ^β-γ^ angle and the O^2G^-N distance are shown along the X and Y axes, respectively, and indicate the probability (calculated as normalized frequency) of conformations with the corresponding distance values.

MD simulations of the Mg-GTP-Ras/RasGAP system showed that the insertion of the Arg finger, Arg789 of RasGAP, into the active site of Ras led to the formation of additional H-bonds between one of the nitrogen atoms of the Arg789 guanidinium group and O^2A^ and O^3G^ atoms of the phosphate chain (Table 1, Fig. 2A). The other nitrogen atom of the Arg789 guanidinium group formed an H-bond with the backbone carbonyl group of Thr785 of RasGAP, whereas the backbone carbonyl of Arg789 interacted with the backbone nitrogen atom of Gln61 forming a weak H-bond with the backbone nitrogen atom of Gln61. Perhaps owing to this interaction, the side chain of Gln61 turned towards the phosphate chain in the Ras/RasGAP complex and made an H-bond with the O^3G^ atom of γ-phosphate (Table 1, Fig. 2A).

In the MD simulation of Ras, Tyr32 was initially in the closed state, forming an H-bond with O^3G^ (Table 1, Fig. S3). This interaction was maintained for the most part of the simulation with the exception of one case, when the bond was broken and Tyr32 moved into the open conformation, but returned to its original position after 2 ns (Fig. S3). In the Mg-GTP-Ras/RasGAP system, Tyr32 was always in the open conformation (Fig. S3); the H-bond of O^3G^ with Tyr32 in Ras was replaced by an H-bond with the Arg789 of RasGAP and Gln61 (Fig. S3).

We also examined the location of water molecules in the active sites of Ras and Ras/RasGAP during MD simulations. In Mg-GTP-Ras, water molecules were mostly seen in two sites near the γ-phosphate (Fig. S4). In the first site, the water molecule was coordinated by O^1G^ and the backbone amide group of Thr35. In the second site, the water molecule was coordinated by O^2G^ and the backbone carbonyl group of Gly60. However, in the absence of RasGAP, this area of protein is exposed, so that water molecules could easily exchange with solution. In the Mg-GTP-Ras/RasGAP complex, only one water molecule remained bound near the γ-phosphate, coordinated by O^1G^, the backbone amide group of Thr35, and the side chain of Gln61. In this case, the active site was closed and the water molecule could not escape the binding site (Table 1, Fig. S4). As compared to Ras, this water molecule was shifted towards the expected in-line attacking position of the catalytic water molecule.

Introduction of the Arg finger between O^2A^ and O^3G^ atoms led to the shortening of the distance between these atoms (Fig S5). In the MD simulation of H-Ras, the average distance between these two oxygens was 5.4 Å, whereas upon the insertion of the Arg residue into the Ras/RasGAP complex, the average distance between them decreased to 4.7 Å (Fig S5). By pulling the O^3G^ atom closer to O^2A^, Arg789 twisted the γ-phosphate, so that its O^2G^ atom formed a novel H-bond with the backbone nitrogen of Gly13 (Table 1, Fig. 2, S3). This bond with Gly13 is not seen in the MD tracks of Mg-GTP-Ras (Table 1) and appears to complement the bond of O^2G^ with Gly60, which has become longer and weaker in the Mg-GTP-Ras/RasGAP complex (Table 1, Fig. S2). The H-bond between Gly13 and O^3B^ was also elongated in the Ras/RasGAP complex, as compared to Ras (Table 1, Fig S2).

To specify the conformational changes of the phosphate chain in response to the formation of the Ras/RasGAP complex, we measured the values of dihedral angles between phosphate groups, namely Ψ^α-β^ = ∠O^2A^-P^A^-P^B^-O^2B^; Ψ^β-γ^ = ∠O^1B^-P^B^-P^G^-O^1G^; and Ψ^α-γ^ = ∠O^1A^-P^A^-P^G^-O^3G^ (Fig. 3, S2). Values of all dihedral angles, as averaged from MD simulations and measured from crystal structures, are summarized in Table S2. In our simulations, the α-phosphate oxygens remained close to the energetically favorable staggered conformation relative to the β-phosphate oxygen atoms in both MD simulation systems, similarly to the crystal structures shown in Fig. 1B and 1C; still, the Ψ^α-β^ angle decreased slightly from 59° to 46° (Fig. 3A). Owing to the coordination of β- and γ-phosphates by the Mg^2+^ ion and Lys16, the dihedral angle between β- and γ-phosphates was close to the eclipsed value of zero degrees both in the crystal structures of H-Ras and Ras/RasGAP (Fig. 1B) and in MD simulations (Fig. 3B, Table S2). In the case of H-Ras, the dihedral angle between α- and γ-phosphates was close to that for the staggered conformation (~70°, Fig. 3C, Table S2). However, in the MD simulation of the Ras/RasGAP complex, γ-phosphate rotated by ca. 40° and approached the eclipsed conformation relative to α-phosphate with Ψ^α-γ^ =31° (Fig. 3C, Table S2). Thus, in the Ras/RasGAP complex, the Arg finger brought the phosphate chain into a state where all three phosphate groups were almost eclipsed (Fig. 2A, 3A-C). While the Arg finger made H-bonds with O^2A^ and O^3G^, the eclipsed conformation was achieved largely due to the rotation of γ-phosphate, accompanied by minor movements of α- and β-phosphates (Fig. 2, 3, S3, Table S2). Although a minor displacement of α-phosphate was observed in the MD simulation (Fig. 2), the H-bond between O^1A^ and the backbone nitrogen of Ala18 was maintained both in Ras and Ras/RasGAP structures (Fig. S3, Table 1). In contrast, γ-phosphate rotated from a near-eclipsed into a perfectly eclipsed conformation with respect to β-phosphate and from a perfectly staggered to a near-eclipsed conformation with respect to α-phosphate (Fig. 2, 3, Table S2). This rotation was accompanied by formation of a new H-bond with Gly13 (Fig. 2, S3, Table 1).

**Figure 3.**
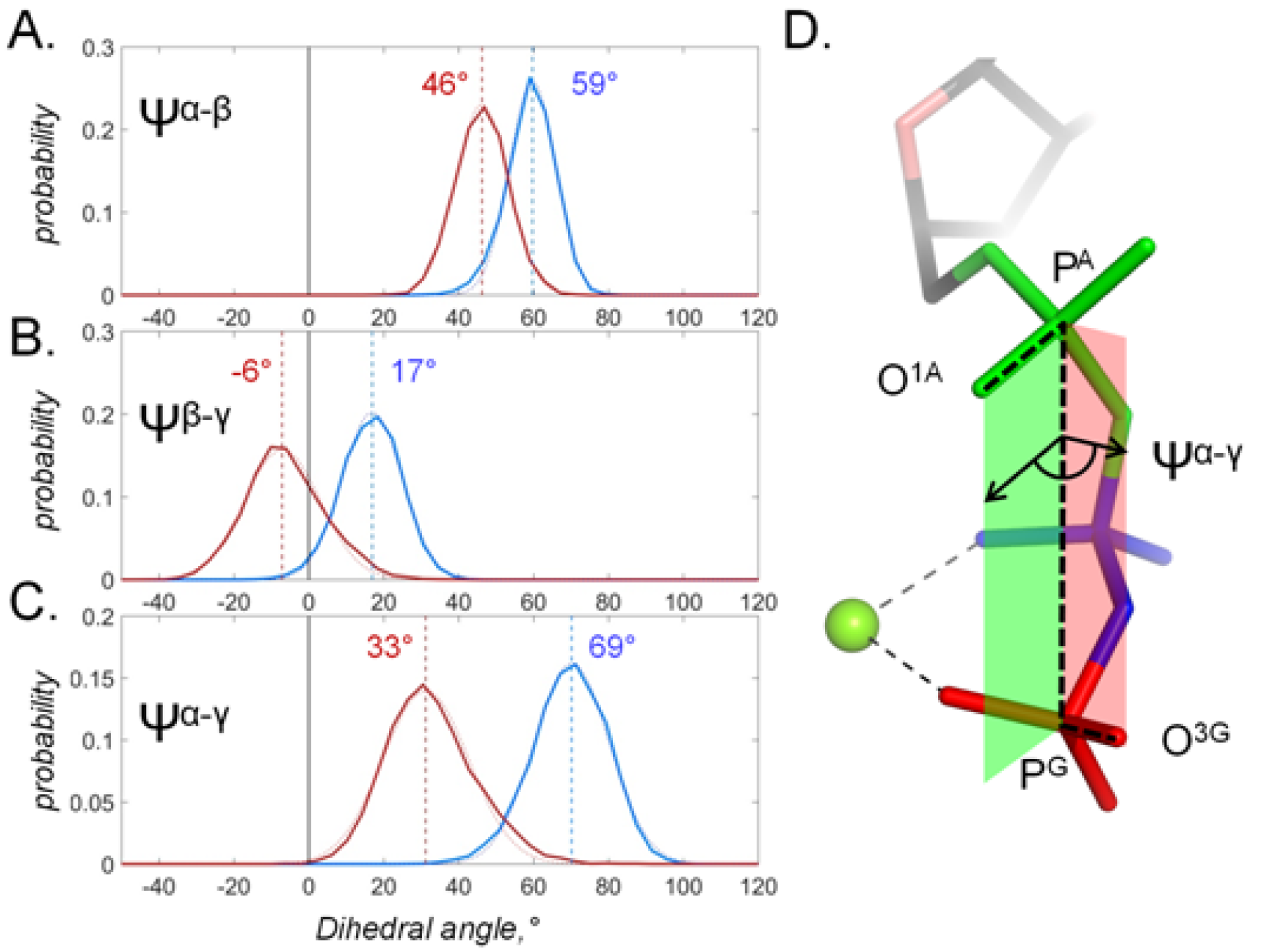
Dihedral angles in GTP molecules bound to Ras and Ras/RasGAP. **A-C.** Distribution histograms for dihedral angles between phosphate groups in GTP, calculated from MD simulations of Mg-GTP-Ras (blue) and Mg-GTP-RasGAP (red). Normalized histograms of dihedral angle distribution (solid lines) were calculated from MD trajectories and fitted with normal distribution function (dotted lines). Red and blue vertical lines indicate the centroid values of the fits by Gaussian function. Black vertical lines indicate Ψ =0°, which corresponds to the fully eclipsed conformation, while Ψ=±60° corresponds to the fully staggered conformation. **D.** The phosphate chain of GTP, colored as in Fig. 1A, illustrating the dihedral angle Ψ^α-γ^. Dihedral angle is an angle between two planes and is defined by 4 atoms. In this case, the angle Ψ^α-γ^ is an angle between the plane that contains atoms P^G^, P^A^ and O^1A^ (green), and the plane that contains atoms P^A^, P^G^ and O^3G^ (red). In the fully eclipsed conformation, both P-O bonds are coplanar, so that the two planes overlap and the dihedral angle between them is 0°.

To estimate the difference in the free energy between the staggered (Ras) and near eclipsed (Ras/RasGAP) states, we calculated the free energy profiles of the γ-phosphate rotation relative to α-phosphate and β-phosphate groups, respectively (Fig. S6). The transition towards the eclipsed conformation in the Ras/RasGAP complex, accompanied by the rotation of γ-phosphate by ca. 40°, corresponded to a change in the free energy of 200-250 meV or ~20-25 kJ/mol.

### Catalytically important residues in the Ras-like GTPases and their complexes with GAP domains

Comparative structural analysis of Ras-like proteins (Fig. 1B-D, Table S3), as well as comparison of the Mg-GTP-Ras and Mg-GTP-Ras/RasGAP structures after their MD equilibration (Fig. 2, 3, S3, Table 1), in accordance with available literature data (e.g. (Jin, Molt, et al., 2017; Kamerlin, et al., 2013; Scheffzek, et al., 1997; Schweins et al., 1994; Wittinghofer & Vetter, 2011)), points to the following specific amino acid residues that appear to be important for the catalytic mechanism of Ras/RasGAP (for simplicity, the amino acid numeration of Ras and RasGAP is used):

1. The residues of the P-loop region (Gly13, Val14, Gly15, Lys16, Ser17, Ala18), as well as Gly60, stabilize the phosphate chain and participate in coordination of the Mg^2+^ ion. The Mg^2+^ ion is additionally coordinated by Thr35 and, via a water molecule, by Asp57 of the Walker B motif.
2. Arg789 residue of RasGAP is inserted into the AG site, provides an additional stabilizing positive charge, forms several H-bonds, including bonds with O^2A^ and O^3G^, and appears to twist the γ-phosphate group.
3. The backbone carbonyl of Arg789 forms an H-bond with backbone nitrogen atom of Gln61. The side chain of Gln61 turns and, together with Thr35, stabilizes the catalytic water molecule close to the attack position.

To assess whether this framework of catalytic interactions could be applicable to other P-loop NTPases, we have traced the conservation of the above-described elements within the entire P-loop superfamily.

### Diversity of activation mechanisms in P-loop NTPases

The abundance and diversity of the P-loop NTPase domains with solved 3D structures makes their classification a complex and arduous task. In the CATH database, the P-loop superfamily 3.40.50.300 includes 140 structural clusters and 2,438 functional families (Sillitoe et al., 2015). In the InterPro database (Finn et al., 2017), the “homologous superfamily” IPR027417 has 698 family and domain entries. In the ECOD database (Schaeffer et al., 2017), the topology-level P-loop_NTPase entry contains 193 families. In the Pfam database (Finn et al., 2016), the P-loop NTPase clan CL0023 contains 217 families. Finally, in the SCOP database (Murzin et al., 1995), the P-loop fold entry contains 24 families. However, neither of these databases provides a hierarchical grouping of all these families. Therefore, in our analysis, we used the classification of Leipe and co-authors, which divides P-loop containing NTPases into two divisions with a total of 8 classes with distinctly different structural organization (Leipe et al., 2003; Leipe, et al., 2002). According to this classification, P-loop NTPases form two divisions: the Kinase-GTPase division and Additional Strand Catalytic Glu (E) (ASCE) division (Leipe, et al., 2003; Leipe, et al., 2002), see the Scheme 1.

**Scheme 1.**
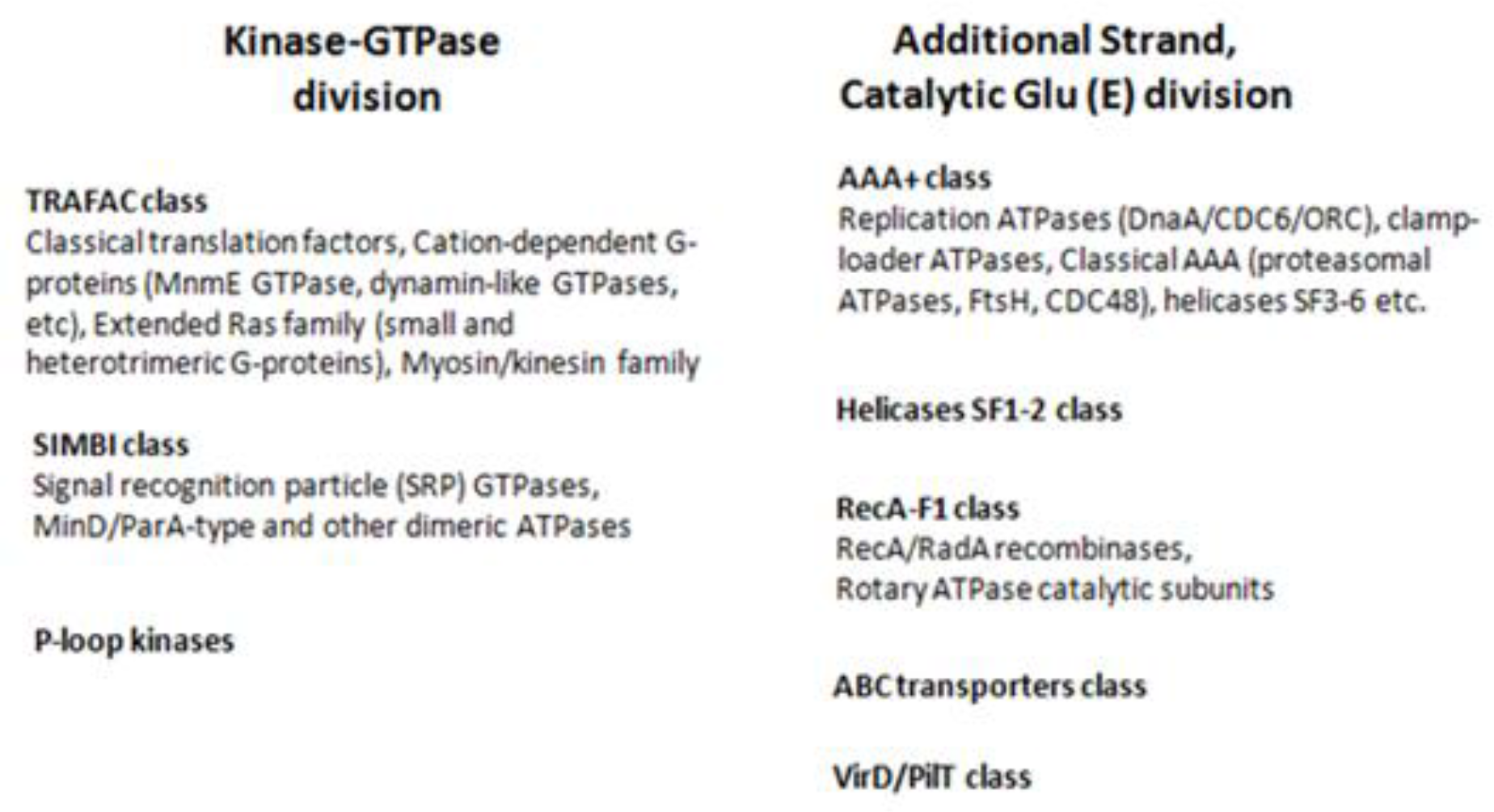
Classification of proteins with the P-loop NTPase fold, as based on refs. (Iyer et al., 2004; Leipe, et al., 2003; Leipe, et al., 2002)).

The Kinase-GTPase division unites 3 classes: TRAFAC class of translational factors and regulatory GTPases, SIMBI class of regulatory dimeric ATPases and GTPases, and the class of P-loop kinases. The ASCE division includes the following 5 classes: AAA+ (ATPases Associated with diverse cellular Activities); RecA-like class, which includes catalytic subunits of F- and V-type rotary ATP synthases; helicases of superfamilies 1 and 2; VirD/PilT ATPases of type II/IV secretion systems, and the ABC transporters (Leipe, et al., 2002). We have limited our structure analysis to these 8 well-studies traditional classes.

We have selected from one to three proteins with available crystal structures to represent each class, based on the available data on the catalytic mechanisms and structural features for each class. Each selected protein structure was superposed with the structure of Ras/RasGAP [PDB 1WQ1] by aligning 27 residues of the P-loop region (residues 2-28 of Ras). Based on the location in the structure and the literature data, we identified residues that correspond to the catalytically important residues of Ras, as described above. We have searched for the following residues in each representative protein: (i) the Arg finger or any other moiety in the AG site; (ii) residues that coordinate the Mg-triphosphate moiety, and (iii) residues in the proximity of γ-phosphate that could be involved in stabilization of the attacking water molecule and are therefore characterized as catalytic/essential in the literature. The results of our analysis are given below and summarized in Table 2 and Fig. 4-6.

**Table 2.**
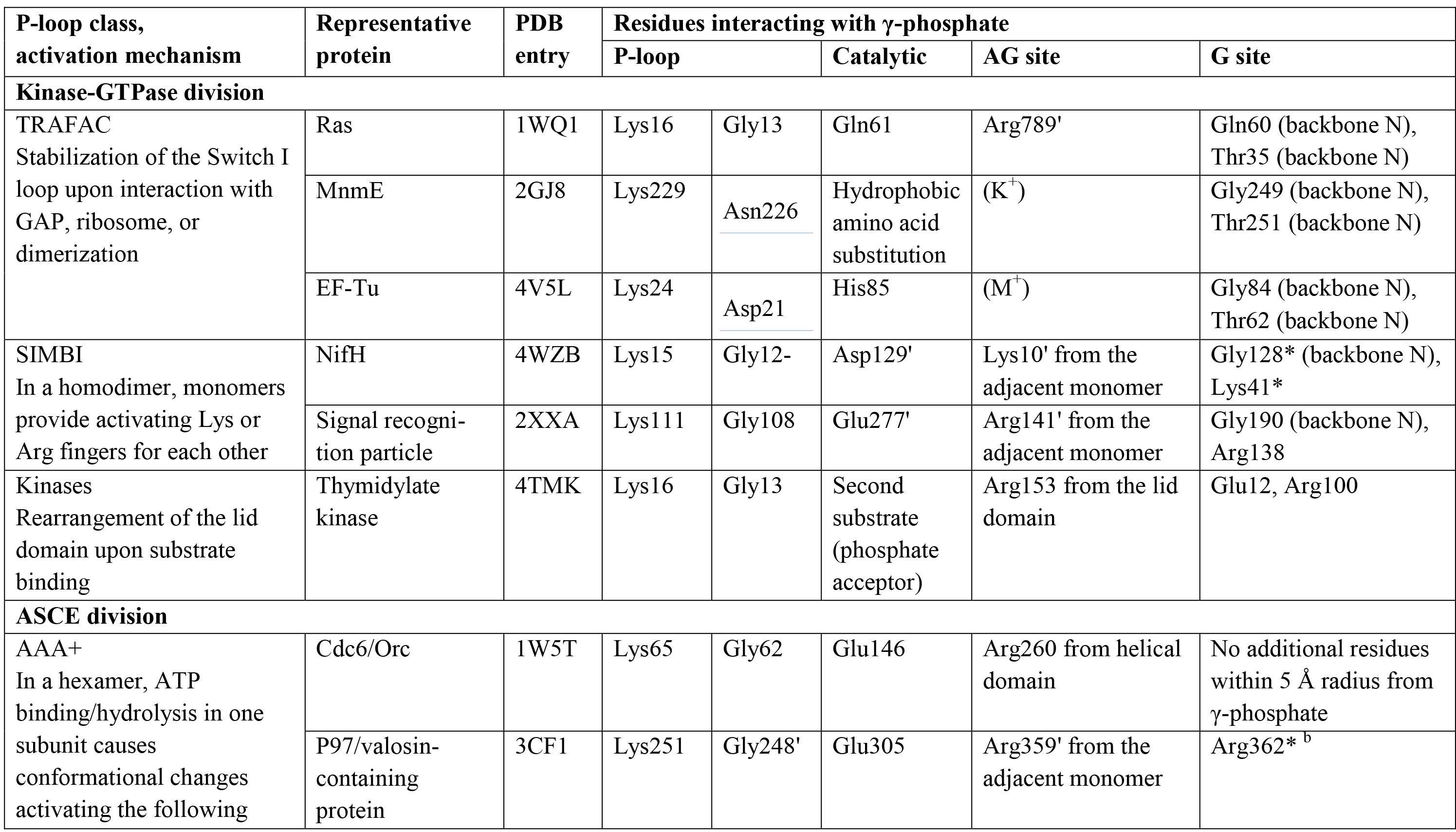
Activation mechanisms in different classes of P-loop NTPases

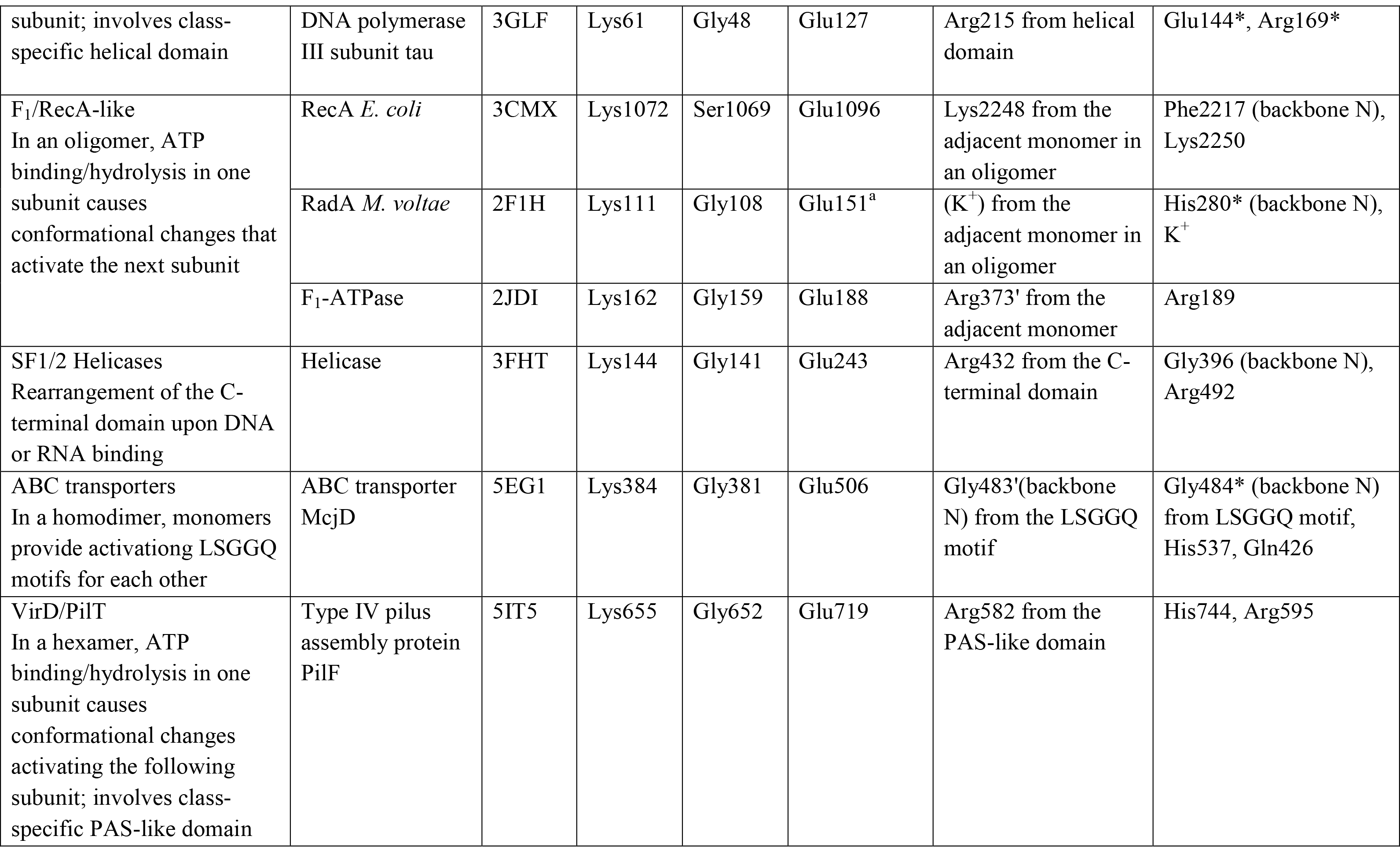

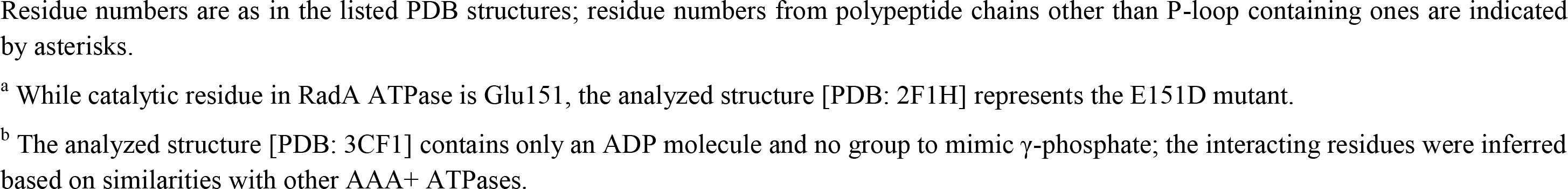

**Figure 4.**
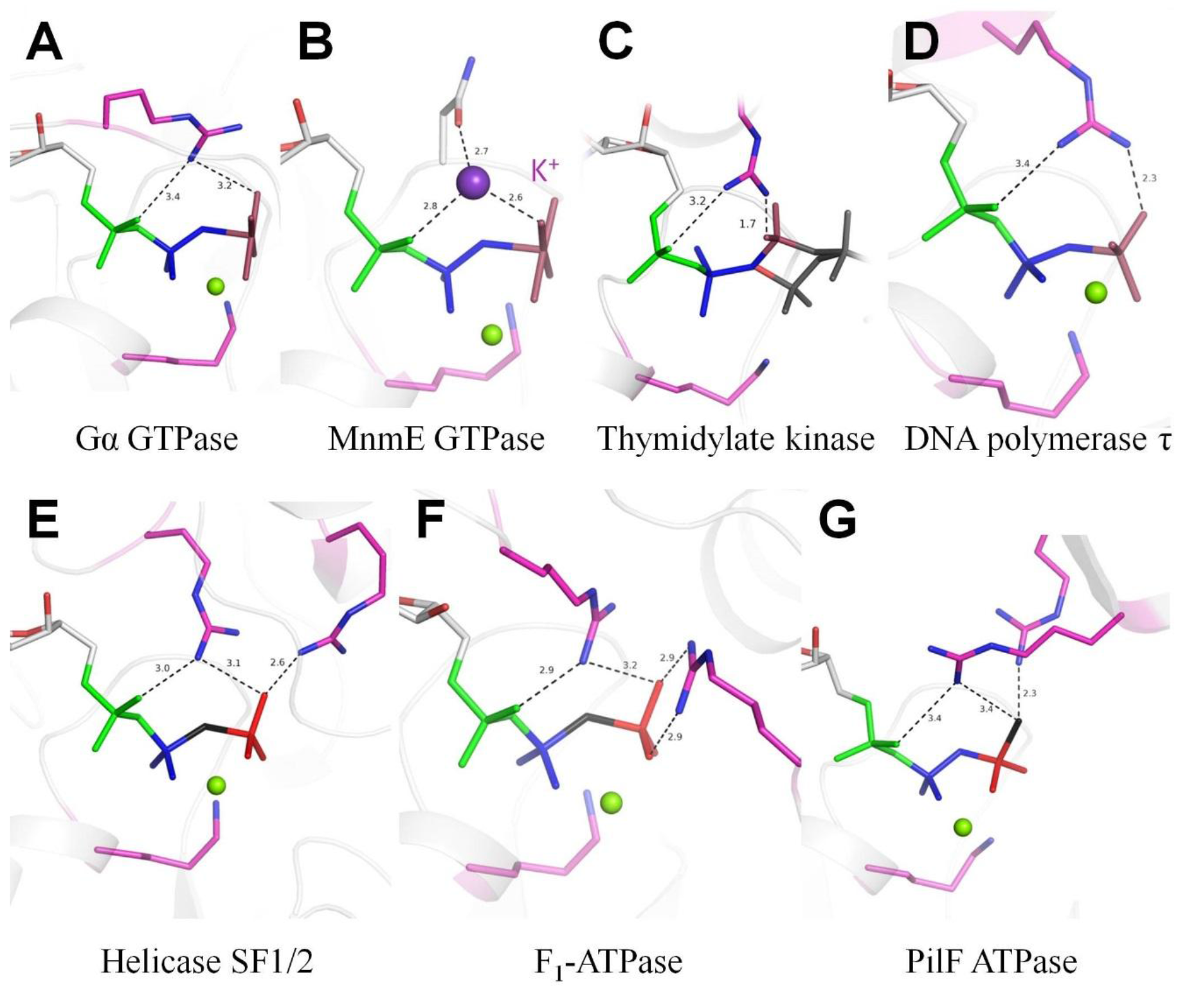
Crystal structures of the catalytic sites of P-loop NTPases with the activating moiety interacting with α- and γ-phosphates. Protein backbones are shown as grey cartoons, nucleotides and their analogs are shown as sticks, phosphate chain is colored as in Fig. 1A. Atoms that are substituted in non-hydrolyzable analogs are shown in black; γ-phosphate mimicking groups in transition state analogs are shown in pink; Lys and Arg residues are shown in magenta; Mg^2+^ ions are shown as green spheres, K^+^ ion is shown as a purple sphere. In panel C, the second substrate-mimicking kinase inhibitor is shown as grey sticks. Distances are in Å. **A**, Heterotrimeric GTPase subunit Gα12 in complex with GDP, Mg^2+^ and AlF_4_^−^ [PDB: 1ZCA]; **B**, Cation-dependent GTPase MnmE in complex with GDP, AlF_4_^−^, Mg^2+^ and K^+^ [PDB:2GJ8]; **C**, Thymidylate kinase in complex with the bisubstrate inhibitor Tp5a [PDB: 4TMK]; **D**, DNA polymerase III subunit tau of the *E. coli* clamp loader in complex with ADP, BeF_3_ and Mg^2+^ [PDB: 3GLF]; **E**, ATP-dependent RNA helicase DDX19B in complex with AMP-PNP and Mg^2+^ [PDB: 3FHT]; **F**, F_1_-ATPase with AMP-PNP bound to the β subunit [PDB: 2JDI]; **G**, Type IV pilus assembly ATPase in complex with ATPγS and Mg^2+^ [PDB: 5IT5].

#### Kinase-GTPase division

##### Class 1. TRAFAC

Members of the TRAFAC class show the broadest variety of activating interactions. This class unites proteins that can be activated either by an interaction with a specific protein (e.g. Ras-like proteins (Bos et al., 2007; Cherfils & Zeghouf, 2013)), or by dimerization (MnmE, dynamins (Chappie et al., 2010; Gasper et al., 2009)), or by interaction with the ribosomal RNA and proteins (classical translation factors (Rodnina et al., 2000) and other ribosome-associated GTPases (Caldon et al., 2001)). Activation of proteins in the TRAFAC class is governed by movement of two loops, following the P-loop: switch I and switch II. These regions undergo conformational changes upon binding of the substrate and activation of the hydrolysis reaction by interaction partners (which could be a specific protein, the second monomer in a dimer, or RNA or DNA).

Our structural analysis showed that the following TRAFAC class protein families mostly rely on the use of Arg fingers: GB1/RHD3-type GTPases (e.g. atlastin), the extended Ras-like family (including heterotrimeric G-proteins), and a deviant group of septin-like proteins within the TEES (TrmE-Era-EngA-EngB-Septin-like) family. The Arg finger can be provided by another protein (as in Ras/RasGAP complex), by another domain of the same protein (e.g. Gα of heterotrimeric GTPases, Fig. 4A of the main text), or by another subunit in an oligomer (e.g. atlastin, septin, hGBP).

In addition, TRAFAC class enzymes can use monovalent cations (M^+^) as activating moieties (Ash, et al., 2012). Families with established cation-dependent activation include classical translation factors, OBG-HflX-like family, TEES, YlqF/YawG-like, and dynamin-like families. In these families, cation binding requires a fixed position of the Switch I loop (dubbed K-loop for K-dependent G-proteins) which, as discussed in detail in (Shalaeva et al., submitted), is governed either by a specific activating interaction of the P-loop domain with another subunit in a dimer (MnmE GTPase, dynamins) or by an interaction with the ribosome (OBG-like proteins, possibly classical translation factors). In most families of M^+^-dependent NTPases of the TRAFAC class, hydrolysis is activated by K^+^ ions, as e.g. in the GTPase MnmE (Fig. 4B of the main text). In the unique case of dynamins, hydrolysis can be activated by either K^+^ or Na^+^ ions (Fig. 5A of the main text). While the activating K^+^ ion or an Arg finger in the AG site interacts with both α- and γ-phosphates (Fig. 4B of the main text), the smaller Na^+^ ion in the same position cannot reach both these phosphate groups at the same time. Instead, as shown in Fig. 5A of the main text, the Na^+^ ion interacts only with the γ-phosphate group, see (Shalaeva et al., submitted) for details.

**Figure 5.**
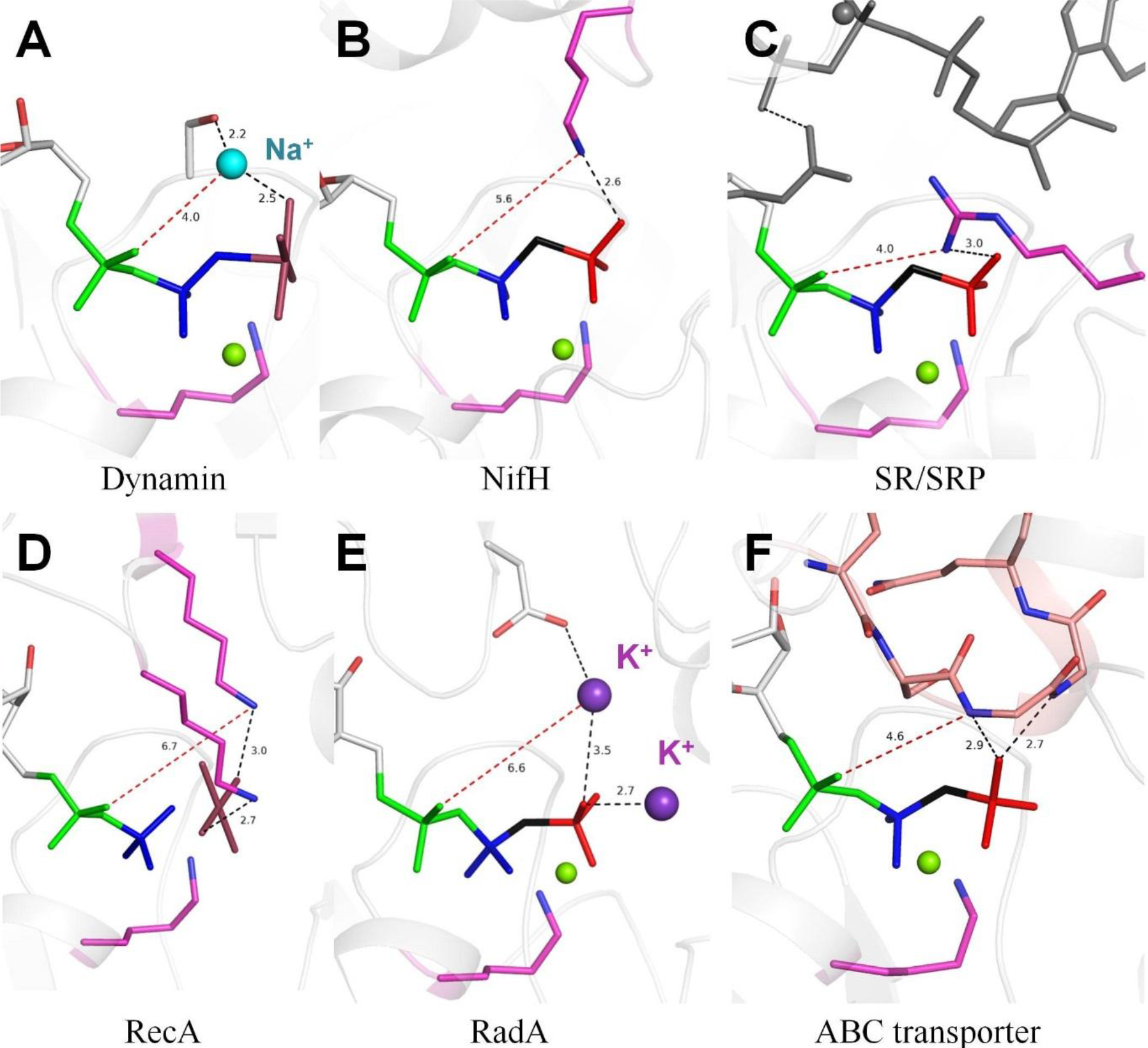
Crystal structures of the catalytic sites of P-loop NTPases with the activating moiety interacting with γ-phosphate. Protein backbones, nucleotides, phosphate chain, and Mg^2+^ and K^+^ ions are colored as in Fig. 4, the Na^+^ ion in panel A is shown as a light blue sphere. Black dashed lines indicate H-bonds of less than 3.5 Å, red dashed lines indicate longer distances. All distances are in Å. **A**, Dynamin GTPase in complex with GDP, AlF_4_^−^, Mg^2+^, and Na^+^ [PDB:2X2E]; **B**, Nitrogenase iron protein 1 (NifH) in complex with AMP-PCP and Mg^2+^ [PDB: 4WZB]; **C**, Signal recognition particle (SRP) bound to its receptor (SR) in complex with GMP-PCP and Mg^2+^ [PDB:2XXA], substrate and the Arg finger in the complementary binding site are shown as grey sticks; **D**, recombination protein RecA oligomer in complex with ADP, AlF_4_^−^ and Mg^2+^ [PDB: 3CMX]; **E**, Archaeal recombinase RadA in complex with AMP-PNP, Mg^2+^ and K^+^ [PDB: 2F1H], adjacent monomer reconstructed from superposition with [PDB: 3CMX]; **F**, ABC transporter McjD in complex with AMP-PNP and Mg^2+^ [PDB: 5EG1].

Highly specialized motor proteins of myosin and kinesin families also belong to the TRAFAC class. Being structurally and functionally highly derived, they use intrinsic Arg fingers, which are positioned at the distal end of the phosphate chain, close to the P^G^ atom.

In addition to cation-binding residue(s) or Arg fingers, NTP hydrolysis in TRAFAC proteins requires the presence of a catalytic water molecule, usually coordinated by a conserved residue from Switch II, see (Jin, Molt, et al., 2017; Jin, Richards, et al., 2017; Kamerlin, et al., 2013; Wittinghofer & Vetter, 2011) for reviews. In the cases of Ras-like GTPases and the Gα subunits of heterotrimeric G-proteins, a highly conserved Gln residue (corresponding to Gln61 in Ras) is thought to be crucial for catalysis. This residue is brought to the active site only upon interaction with GAP (in Ras-like proteins) or helical domain (in heterotrimeric G-proteins). In translation factors, this role is played by a His residue in the same position in Switch II, but orientation of this residue towards γ-phosphate is triggered by the interaction of the protein with the ribosome. Finally, a subset of GTPases termed HAS-GTPases possess a hydrophobic residue in the position of the catalytic Glu residue (hence HAS – hydrophobic amino acid substitution). The Thr residue of Switch I, which is involved in Mg^2+^ coordination sphere and is universally conserved in the TRAFAC class, can also provide its backbone oxygen atom to coordinate the catalytic water molecule.

The family of Ras-like proteins can be divided into five principal subfamilies: Ras, Rho, Rab, Arf, and Ran (Wennerberg et al., 2005). In most cases, GAPs provide an Arg finger to the active site of Ras-like GTPase and stabilize the position of the catalytic Gln residue (Gln61 in Ras), see (Bos, et al., 2007; Cherfils & Zeghouf, 2013; Jin, Richards, et al., 2017; Kamerlin, et al., 2013; Ligeti et al., 2012; Sprang, 1997; Wittinghofer & Vetter, 2011) for reviews. This canonical activation mechanism has been established for Ras and Rho GTPases. Similarly, Arg finger from GAP is employed in Arf GTPases, although the catalytic Gln is replaced there with His (Renault et al., 2003). In Rab GTPases, GAPs provide both an Arg finger to the AG site and the catalytic Gln residue (Pan et al., 2006).

While most Ras-like GTPases employ an Arg finger, two exceptions appear to exist, namely Rap and Ran GTPases. The Rap GTPase lacks the catalytic Gln61, which is replaced with Thr (Scrima et al., 2008). Initially described RapGAPs have different fold than RasGAPs and catalyze the GTP hydrolysis by providing an Asn residue (referred to as the ‘Asn thumb’) (Scrima, et al., 2008). GTP hydrolysis in Rap, however, can also be activated by plexins and dual-specificity GAPs, which share the RasGAP fold and provide a classical Arg finger (Kupzig et al., 2006; Pena et al., 2008; Y. X. Wang et al., 2013). The structure [PDB: 1K5D] of Ran-RanGAP complex places Tyr39 of RanGAP into the AG site instead of the Arg finger. However, the side chain of Arg170 of RanGAP is located near the active site and potentially could reach the AG site. In the crystal structure, this residue is involved in a salt bridge with Glu172 of RanGAP (Seewald et al., 2002). Since Tyr39 cannot provide a positive charge needed for activation, it could not be excluded that Arg170 serves as the activating finger *in vivo*.

##### Class 2. SIMBI

P-loop NTPase class SIMBI includes ATPases and GTPases which get activated by dimerization, upon which the active sites of monomers interact with each other. In such dimers, monomers provide to each other’s active sites either activating Lys residues (Fig. 5B of the main text), or Arg residues (in the case of signal recognition particles and their respective receptors that are members of the SR/SRP-like families, see Fig 5C of the main text) (Bange & Sinning, 2013; Lutkenhaus, 2012).

In the SRP/SR and FtsY/Ffh complexes, the mutual exchange of Arg fingers between monomers is not sufficient to trigger the hydrolysis reaction. Only upon interaction of such dimer with the RNA tetraloop, the catalytic Glu/Gln residue of SRP (Glu277 in PDB entry 2XXA) is turned towards the γ-phosphate group of the SR-bound GTP (Ataide et al., 2011).

Typical SIMBI ATPases and GTPases contain the so-called ‘deviant Walker A motif KGGXXGKT’ with an additional conserved Lys that is introduced to the active site of the second subunit upon dimerization (Lutkenhaus, 2012). Nucleotide-dependent dimerization in such proteins is associated with conformational changes in Switch I and Switch II regions, similarly to the activation of TRAFAC GTPases. Switch II contains two residues (Asp166 and Thr167 in PDB entry 2WOJ) which make second-shell interactions with Mg^2+^ ion. Switch I region contains a conserved Asp residue (Asp57 in PDB 2WOJ), which coordinates the water molecule positioned for in-line nucleophilic attack on the γ-phosphate of the substrate (Mateja et al., 2009).

##### Class 3. Kinases

P-loop kinases are ubiquitous enzymes that transfer the γ-phosphoryl residue of ATP to a wide range of substrates, primarily nucleotides and other small molecules(Cheek et al., 2005; Leipe, et al., 2003). The key role in the catalysis by P-loop kinases is played by structurally conserved Arg residue located in the so-called “lid” domain, a small helical bundle domain that often carries several Arg residues (Cheek, et al., 2005; Kenyon et al., 2012; Leipe, et al., 2003), see Fig. 4C of the main text.

P-loop kinases do not need other proteins or domains to provide activating Arg/Lys fingers or to induce conformational changes in any “switch” regions. Instead, binding of the substrate nucleotide is sufficient to trigger the lid domain rearrangement that results in the introduction of Arg finger(s). The ATP molecule itself mediates deprotonation of the–OH group of the phosphate acceptor molecule which, instead of the water molecule in hydrolysis reactions, initiates the nucleophilic attack and formation of the pentavalent intermediate to allow phosphoryl transfer (Kenyon, et al., 2012).

#### ASCE division

##### Class 4. AAA+ proteins

AAA+ proteins are ATPases associated with various cellular activities, where “+” stands for ‘extended’. These enzymes carry an additional α-helical C-terminal domain with conserved Arg residues that are present in most families (Ogura, et al., 2004). At the same time, the core P-loop domain of the AAA+ ATPases carries another conserved Arg residue, usually referred to as “Arg finger” and characterized as crucial for catalysis (Karata et al., 1999). P-loop domains of AAA+ proteins interact upon oligomerization (most often a hexamer is formed (Wendler, et al., 2012)), so that the nucleotide binding site of one subunit receives an Arg finger from the neighboring subunit and, in some protein families, an additional Arg finger from its own helical domain (Fig. 4D of the main text). Oligomerization of AAA+ proteins leads to conformational changes in the Sensor I loop of the P-loop domain that carries Thr and Asn residues (Wendler, et al., 2012) and, together with the signature catalytic Glu, forms a network of interactions between two subunits. This network includes the catalytic water molecule Sensor 2 loop, which is located on the opposite site of the protein globule relative to the P-loop and contains Arg residue(s) introduced into the active site of the neighboring subunit, as well as, in some protein families, the Arg finger of the helical domain that is also introduced into the active site (Wendler, et al., 2012). Thus, the active site of each monomer in an AAA+ oligomer receives an Arg residue from Sensor 2 of the P-loop domain of another monomer and often, but not always, the second Arg residue that is provided by the helical domain of the same monomer. When two positively charged residues are present in the nucleotide-binding site, one of them occupies the AG site of the substrate phosphate chain, while the other one is located at the end of the phosphate chain in the proximity of γ-phosphate(Wendler, et al., 2012).

##### Class 5. Helicases of subfamilies 1 and 2

Helicases are a large group of proteins with the ability to unwind DNA and/or RNA helices. Based on the sequence and structural similarity, helicases are traditionally divided into six superfamilies (SF1-SF6) (Singleton et al., 2007). NTP-binding domains of all helicase superfamilies belong to the P-loop fold, with helicases from superfamilies SF4, SF5, and SF6 usually attributed to the AAA+ class (Ammelburg et al., 2006). Helicases of superfamily SF3 also have AAA+-like P-loop domains but are excluded from that class because of an aberrant topology of the C-terminal helical domain (Ammelburg, et al., 2006). Helicases of superfamilies 1 and 2 (SF1/2) form a separate class among the ASCE ATPases. While helicases from SF3-SF6 and many other AAA+ family proteins typically act as hexamers, SF1/2 helicases are mostly monomeric or dimeric but each polypeptide chain of these proteins contains two domains with the P-loop fold (Singleton, et al., 2007). In such helicases, the P-loop of N-terminal domain carries functional Walker A and B motifs and binds the nucleotide molecule, while the C-terminal domain, despite also having a P-loop-like fold, has lost those motifs and does not bind NTPs. Upon interaction with the RNA or DNA helix, the C-terminal domain provides two Arg residues to the nucleotide-binding site of the N-terminal domain: one into the AG site and one to the end of the phosphate chain to bind γ-phosphate, see Fig. 4E of the main text and (Sengoku et al., 2006).

The SF1 and SF2 helicases convert the chemical energy of ATP hydrolysis into the mechanical energy required for DNA unwinding and translocation. Precisely how this coupling is achieved appears to be different between particular SF1 and SF2 enzymes. However, structural rearrangement caused by DNA or RNA binding seems to not only introduce Arg fingers via inter-domain interactions, but also couple family-specific substrate binding with the movement of the invariable catalytic Glu residue (Bhattacharyya & Keck, 2014).

##### Class 6. VirD/PilT-like

Type II/IV secretion system (T2SS) ATPases and related proteins form a separate class of ASCE proteins, most known for its members PilT, PilB, GspE and VirD. These proteins form hexamers which are incorporated into multi-subunit membrane-associated protein secretion system apparatus (Korotkov et al., 2012). In addition to the nucleotide-binding P-loop domain, which is similar to the one in other ASCE ATPases, T2SS ATPases contain an N-terminal two-layer α/β sandwich domain, similar to the well-known ligand-binding PAS domain (Ponting & Aravind, 1997). In the hexamer, the PAS-like domain provides the Arg finger to the AG site of the NTP in the same subunit, see Fig. 4G of the main text and (Mancl et al., 2016; Satyshur et al., 2007). Similarly to other ASCE ATPases, the signature Glu residue is observed in the proximity of γ-phosphate and is expected to polarize the catalytic water molecule (Mancl, et al., 2016).

##### Class 7. RecA-like

This class encompasses oligomeric ATP-dependent motors, involved in homologous recombination and DNA repair (RecA family), as well as α/β subunits of rotary F- and V-type ATP synthases. These proteins do not involve additional domains to provide an Arg finger. Instead, an Arg residue (in rotary ATPases, see Fig. 4F of the main text) or two Lys residues (in recombinases, see Fig. 5E of the main text) are provided by the P-loop domain of the neighboring subunit in the hexamer (or other homooligomer).

Similarly to other proteins from the ASCE division, RecA-like recombinases recruit a Glu residue to coordinate the catalytic water molecule. However, while previously mentioned classes (AAA+, SF1/2 helicases, and VirD/PilT) use the catalytic Glu that directly follows the Walker B motif in the sequence, in RecA-like recombinases and rotary ATPases, the conserved Glu responsible for coordination of the catalytic water molecule is positioned on another strand (Yamaichi & Niki, 2000). This strand is located in the β-sheet of the P-loop domain between the strands that carry the Walker A and the Walker B motifs.

##### Class 8. ABC transporters

The ATP-binding domains of ABC (ATP-binding cassette) family transporters are the only members of ASCE-type NTPases that function as homodimers. In the same way as in SIMBI NTPase dimers, nucleotide-binding sites of ABC transporters are on the interface between the dimer subunits. Instead of an Arg or Lys residue, each monomer inserts the whole signature motif LSGGQ into the catalytic pocket of the other monomer. The backbone amide groups of the two sequential glycine residues occupy the space between the α- and γ-phosphates of the ATP phosphate chain and, by forming H-bonds with γ-phosphate, communicate the changes in the active site to the neighboring subunit, see Fig. 5F and (Chen et al., 2003; Hopfner et al., 2000; Jones & George, 1999).

Several residues commonly found in the active sites of ABC transporters could potentially stabilize the attacking water molecule. This includes Glu residue adjacent to the Walker B motif (similarly to other classes of ASCE proteins), Gln residue(s) from the Q-loop on the additional strand, and a His residue in the H-motif region, which is specific for the ABC transporters (Rees et al., 2009). So far, the residue serving as the (primary) catalytic residue remains ambiguous. While there is strong evidence in some systems that the Glu adjacent to the Walker B motif is the crucial catalytic residue, this may not be true for all proteins within the class (Jones & George, 1999).

#### Comparative structure analysis of activation mechanisms in P-loop NTPases

##### Activating moieties in the AG site

Our comparative structural analysis showed that each class of P-loop NTPases can be characterized by its signature mechanism of introducing the activating cationic moiety into the active site in a controlled way, see SI. The mechanism is shared by most, if not all, protein families in each particular class. In most cases, such mechanisms involve interactions of the P-loop domain with another domain or protein. P-loop NTPases could be activated by dimerization (SIMBI class and ABC transporters), oligomerization (RecA-like and AAA+ classes) or interaction with another domain of the same protein (kinases and helicases SF1/2). P-loop NTPases of the TRAFAC class employ a variety of activation mechanisms: interaction with a specific protein (e.g. the GAP in case of Ras-like proteins, dimerization (MnmE, dynamins), and domain rearrangement upon interaction with the ribosome (classical translation factors and other ribosome-associated GTPases). Activation mechanisms of all classes of P-loop NTPases are summarized in Table 2 and reviewed in more detail in the SI.

Arginine fingers seem to be most widespread as activating moieties: they are used in most families of the TRAFAC class, as well as in kinases, AAA+ ATPases, SF1/2 helicases, and PilB/VirD ATPases. ATPases and GTPases of the SIMBI class employ both Arg residues (e.g. in signal recognition particles and their receptors) and Lys residues. In the RecA-like class, Lys fingers are used in the RecA and RadA-like families, whereas the catalytic subunits of rotary ATP synthases use Arg fingers. Additionally, some NTPases of the TRAFAC class and some recombinases of RecA-like class can be activated by K^+^ or NH_4_^+^ ions; the activation, however, requires interactions with other proteins or domains to complete the cation-binding site, see (Shalaeva et al., submitted) for details.

In the Ras/RasGAP complex, the Arg finger interacts with two oxygen atoms, bridging α- and γ-phosphates (Fig. 2); same arrangement is observed in many P-loop NTPases of different classes, as shown in Fig. 4. In several classes of P-loop NTPases, the activating moiety does not interact with α-phosphate, but only with the γ-phosphate group (Fig. 5).

Finally, in the most exotic cases, even small cations (Na^+^ in dynamins, Fig. 5A) or large groups with a partial positive charge (characteristic motif in ABC (ATP-binding cassette) family transporters, Fig. 5F) can be utilized as activating groups. Specifically, upon dimerization of the ABC transporters, each monomer, instead of an Arg or Lys residue, inserts the signature motif LSGGQ into the catalytic pocket of the other monomer (Jones & George, 1999). The backbone amide groups of two sequential glycine residues occupy the AG site of the ATP phosphate chain and provide nitrogen atoms to coordinate the γ-phosphate (Fig 5F). This system with two backbone amide groups, carrying partial positive charges and able to form H-bonds with the oxygen atom of γ-phosphate, is similar to the classical Arg/Lys fingers. However, both Gly backbone amide groups are located too far from the α-phosphate and can only coordinate the γ-phosphate. In addition, βCH_2_-group of Ser residue separates the backbone nitrogen atoms of Gly residues from the free oxygen atom of α-phosphate. Such a construct provides less electrostatic compensation than an Arg/Lys finger or a monovalent cation, but the large size and rigid structure of the activating moiety apparently enables twisting of γ-phosphate.

We have not found a single case where the activating moiety interacted only with the α-phosphate group.

In spite of their diversity, all activating moieties in P-loop NTPases share three common features: (i) presence of a positive charge (even partial positive charges can do, as evident from the ABC transporters); (ii) controlled insertion, i.e. activating groups are introduced into the catalytic site only under certain conditions, and (iii) the ability to interact with the γ-phosphate group.

##### Residues that stabilize the Mg^2+^ ion and triphosphate moiety of NTP

Residues that stabilize the Mg^2+^ ion belong to the P-loop (Walker A) motif (Ser17 in Ras) and the Walker B motif (Asp57 in Ras). While in many classes the conserved Asp residue from the Walker B motif is directly involved in Mg^2+^ coordination, in TRAFAC class this residue is involved in the second coordination sphere, whereas another residue (Thr35 in Ras) not conserved in other classes of P-loop NTPases, is directly coordinating the Mg^2+^ ion. Overall, coordination of the Mg^2+^ ion is very similar among all classes, as it involves several residues that are conserved across all P-loop NTPases.

Most of the residues that stabilize the triphosphate moiety belong to the P-loop motif. Not only the sequence motif, but also the shape of the P-loop is strictly conserved across all classes of P-loop NTPases (Fig 6). Accordingly, the interactions of the P-loop residues with the triphosphate chain should also be conserved across the whole superfamily.

**Figure 6.**
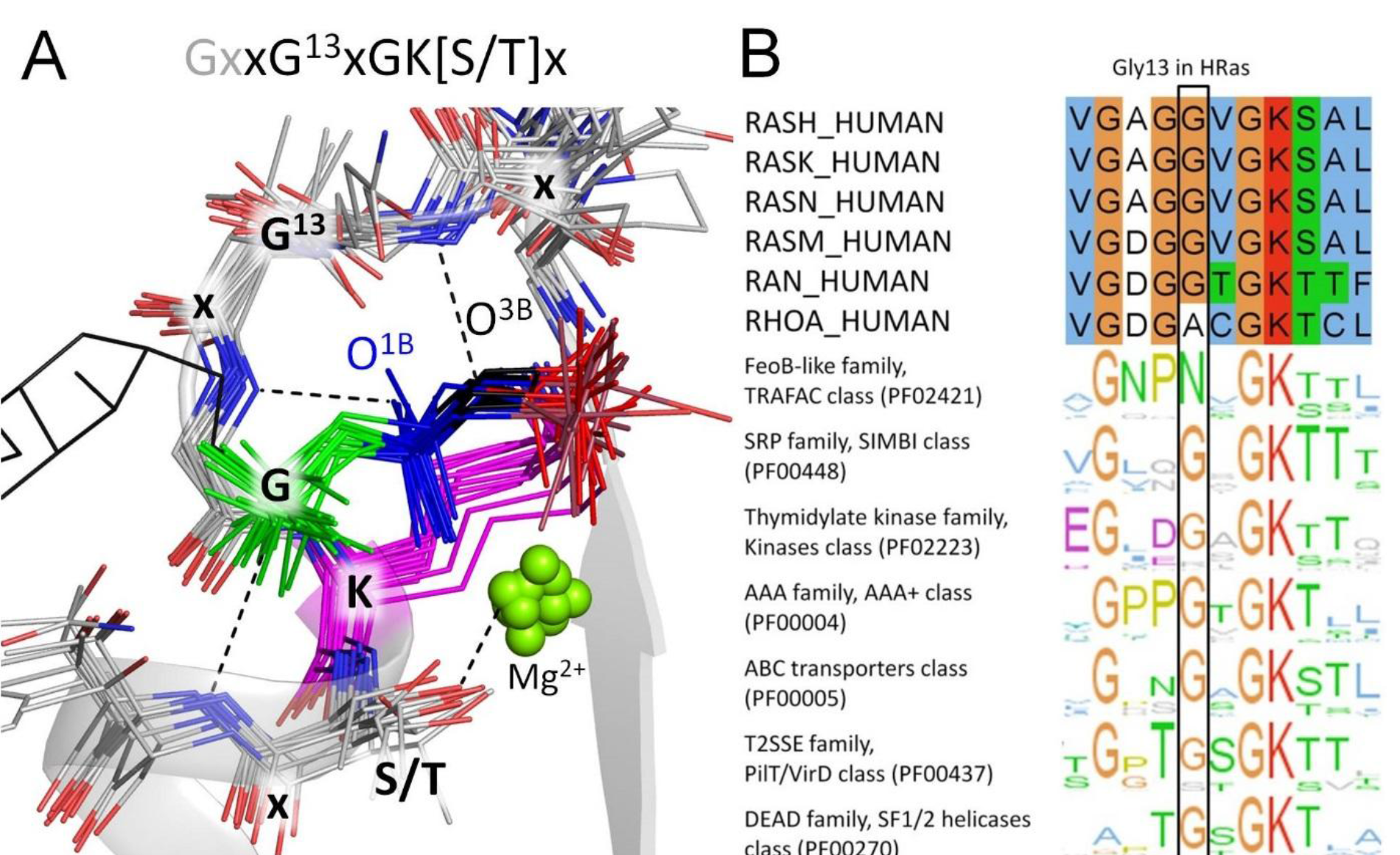
Conservation of the P-loop motif residues across protein families. **A**, Structures of proteins, representing different classes of P-loop NTPases are shown superimposed to the P-loop region of H-Ras [PDB: 1WQ1] (Scheffzek, et al., 1997). For the H-Ras protein, the P-loop region is shown as cartoon, GDP and AlF_3_ are shown as sticks. For all the proteins, the last seven residues from the motif GxxGxGK[S/T]x, as well as phosphate chains and corresponding parts of NTP analogs are shown in sticks and colored as follows: α-phosphate in green, β-phosphate in blue, γ-phosphate in red, the γ-phosphate-mimicking groups in pink; the atoms that are substituted in non-hydrolyzable analogs are colored black. Conserved Lys residues are shown in magenta; Mg^2+^ ions are shown as green spheres. Protein structures are the same as used in Fig. 4 and 5 (PDB IDs: 2GJ8, 2X2E, 2XXA, 4WZB, 2F1H, 3CMX, 2JDI, 5IT5, 1W5T, 3CF1, 3GLF, 3FHT, 5EAG, and 1ZCA). **B**, Conservation of the P-loop sequence motif across protein families. The Sequence Logo representations (Waterhouse et al., 2009) for “seed” alignments of individual protein families from the Pfam database (Finn, et al., 2016) were superposed to match the conserved lysine residue. The box indicates the position that corresponds to Gly13 in human Ras GTPase.

Our analysis showed that other backbone amide groups and/or additional Arg/Lys residues could be involved in coordinating oxygen atoms of γ-phosphate; these residues are indicated for different classes of P-loop NTPases in Table 2 and are poorly conserved. Interaction of additional Arg/Lys residues with the phosphate chain is often promoted by the same protein-protein interaction that provides the activating Arg/Lys finger. For example, the lid domains of kinases, C-terminal domains in SF1/2 helicases, and PAS-like domains in PilT-like proteins often (but not always) carry two Arg residues on the interface directed towards the P-loop domain (see the SI). One Arg residue is inserted into the AG site, while the other one interacts only with γ-phosphate (Fig. 5D, F). In other cases, such as some AAA+ ATPases, Arg finger in the AG site comes from the C-terminal domain of the same subunit and an additional Arg residue next to γ-phosphate is provided by an adjacent subunit (Wendler, et al., 2012), see SI for further details. The positions of these additional partners roughly correspond to the ion-binding site “G“ from the back end of the phosphate chain, as described in (Shalaeva et al. submitted) for the Mg-ATP complex in water. These accessory residues, however, are poorly conserved even within individual families of P-loop NTPases.

##### Residues involved in polarization of the catalytic water molecule

While the activating Lys/Arg fingers or their analogs are usually provided by another protein or domain (Ash, et al., 2012; Jin, Molt, et al., 2017; Kamerlin, et al., 2013; Ogura, et al., 2004; Scheffzek, et al., 1997; Scrima & Wittinghofer, 2006; Wendler, et al., 2012; Wittinghofer & Vetter, 2011), P-loop NTPases generally employ polar residues of their own P-loop domain to assist in coordination and polarization of the catalytic water molecule. Such residues are usually denoted as catalytic or essential, and their replacement by non-polar residues may affect the catalytic function. In the Ras GTPase and related proteins of the TRAFAC class, the catalytic water molecule appears to be coordinated by counterparts of Thr35 and Gln61, see Fig. S4, Table 1 and (Jin, Molt, et al., 2017; Jin, Richards, et al., 2017; Wittinghofer & Vetter, 2011); these residues are not conserved in other classes of P-loop NTPases (Table 2). Furthermore, a family of HAS GTPases in the TRAFAC class lacks a catalytic residue (hence HAS – Hydrophobic Amino acid Substitution) (Mishra et al., 2005). In the ASCE division, the catalytic residue is usually Glu (Table 2), but its exact position varies between classes. In AAA+ ATPases and helicases SF 1/2, the catalytic Glu residue is located on the same β-strand as the Walker B motif, and directly follows the conserved aspartate (Wendler, et al., 2012). In RecA-like recombinases and rotary ATPases, the conserved catalytic Glu is located on an adjacent β-strand from the Walker B motif (Muneyuki et al., 2000). In ABC transporters, the (primary) catalytic residue remains ambiguous (Jones & George, 1999). Finally, in PilT-like ATPases, the catalytic Glu directly follows an atypical Walker B motif, where the Asp residue is replaced with Gly (Mancl, et al., 2016). In the Kinase-GTPase division, not only the position, but also the nature of the catalytic residue varies between classes and families.

## Discussion

The fundamental importance of nucleoside triphosphates, such as ATP and GTP, for biological systems stems from their relative stability and the possibility to dramatically accelerate their hydrolysis by recruiting simple positively charged moieties, such as Mg^2+^ and K^+^ ions and/or side groups of Lys or Arg residues, for catalysis, as well as by using a polarized water molecule to act as a nucleophile (Jin, Molt, et al., 2017; Kamerlin, et al., 2013; Scheffzek, et al., 1997; Schweins, et al., 1994; Wittinghofer & Vetter, 2011).

The abundance of NTP-utilizing enzymes encoded in every living cell makes regulating their activity a key problem in cell metabolism. Obviously, uncontrolled NTP hydrolysis would lead to the depletion of cellular ATP/GTP and be detrimental for cell survival. P-loop ATPases are unique enzymes as they are usually activated before (or during) each turnover. Because of that, the NTP hydrolysis reaction could be divided into two steps. First, an NTP molecule binds to the P-loop domain and attains a catalytically prone conformation; in this conformation hydrolysis proceeds by several orders of magnitude faster than in water; for instance, it is accelerated by five orders of magnitude up to 10^−4^ s^−1^ in case of Ras (Kotting & Gerwert, 2004; Shutes & Der, 2006). Delbaere and coauthors noted that stabilization of β- and γ-phosphates in an eclipsed state by the Mg^2+^ ion and conserved Lys residue of the P-loop (Fig. 1-3) should destabilize the phosphoanhydride bond because of repulsion of oxygen atoms (Delbaere et al., 2004; Matte et al., 1998). The bound NTP molecule, however, still persists on the cellular time scale. The controlled fast hydrolysis takes place upon the interaction with a proper physiological partner, which could be a domain of the same protein, a separate protein or a DNA/RNA molecule (Table 2). Upon this interaction, an activating moiety (an Arg/Lys residue, or a K^+^/Na^+^ ion, or the LSGGQ motif) gets inserted into the binding site (Fig. 4, 5) and promotes the cleavage of the NTP molecule. Still, the exact order of events during the NTP hydrolysis remains obscure. One of the unresolved questions is whether the hydrolysis follows an associative path, being kinetically limited by the nucleophilic attack of a polarized water molecule, or a dissociative path, being limited by the scission of the terminal O^3B^−P^G^ bond (Kamerlin, et al., 2013). Another controversy is whether the proton from the polarized water molecule, on its way to the leaving NDP moiety, is transiently accepted by the catalytic amino acid residue (e.g. Gln61 in case of Ras) (Khrenova et al., 2015) or by the γ-phosphate group (Kamerlin, et al., 2013; Langen et al., 1992; Schweins, et al., 1994).

For the Mg-NTP complex in water, extensive theoretical studies yielded similar barrier heights for the associative and dissociative mechanisms, which prompted suggestions of concerted reaction mechanisms with the nucleophilic attack and bond scission being coupled (Akola & Jones, 2003; Florian & Warshel, 1998; Kamerlin, et al., 2013; Plotnikov et al., 2013; C. Wang et al., 2015). The multiplicity of potential kinetic paths may be the reason why extensive QM/MM calculations, as performed, for instance, for the Ras/RasGAP system (Jin, et al., 2016; Khrenova, et al., 2015; Rudack, et al., 2012), have not yielded consensus results.

These ambiguities prompted us to apply the tools of evolutionary biophysics. With this approach, the key mechanistic elements are identified in well-studied reference systems and then their conservation is checked, from structure and sequence data, throughout the entire protein (super)family. In case of P-loop NTPases, this approach is particularly justified since the experimental investigation of all members of this vast superfamily would not be realistic.

### NTP hydrolysis by the Ras/RasGAP complex as inferred from the MD simulations

Classical MD simulations do not provide direct estimation of kinetic parameters, such as rates and activation energies of chemical reactions. Still, using this approach with H-Ras and Ras/RasGAP enabled valuable sampling of GTP conformations, which is sensitive to details of solvating water structure near the negatively charged triphosphate chain. More sophisticated QM-based methods of dynamics calculation on comparably long time scales are still computationally too expensive for quantitative implementation.

Since crystal structures of the H-Ras/RasGAP complex with a GTP molecule bound are unavailable, there is no full consensus on the exact structure of the H-bond network in the transition state. Our MD simulations yielded an equilibrated structure where one of the nitrogen atoms of the guanidinium group of Arg789 simultaneously formed H-bonds with O^2A^ and O^3G^ in agreement with QM/MM calculations (Klahn et al., 2006), MD simulations (Lu, Jang, Nussinov, et al., 2016; Resat et al., 2001), ^19^F-NMR data obtained with the closely related RhoA/RhoGAP system, see Fig. 1C and (Jin, et al., 2016), numerous crystal structures of GTP analogs containing Ras-like proteins in complex with their activators (Fig. S1C, D), as well as hydrogen bonding patterns of Arg residues as inferred from analysis of PDB structures (Armstrong et al., 2016). In addition, our MD simulations of the H-Ras/RasGAP system showed that the interaction of Arg789 with O^2A^ and O^3G^ results in an almost eclipsed conformation of the phosphate chain, mostly owing to the rotation of the γ-phosphate group (Fig. 2, 3).

The rotation of γ-phosphate is constrained only by its H-bonding with the conserved Lys16 residue of the P-loop, the backbone amide groups of flexible Gly-rich loop regions, and the coordination bond with the Mg^2+^ ion. Owing to the ability of the Lys side chain to stretch out (Cherepanov & Mulkidjanian, 2001), this network of bonds around γ-phosphate appears to be flexible/elastic enough to permit a pronounced rotation of γ-phosphate, bringing it into the perfectly eclipsed conformation relative to the β-phosphate group and to an almost eclipsed conformation relative to α-phosphate (Fig. 2, 3). In this conformation, the repulsion by the negatively charged oxygen atoms of α- and β-phosphates (Cannon, 1993) should push away the γ-phosphate group.

The rotational capacity of γ-phosphate follows already from the comparison of crystal structures (Fig. 1B-D, S1A-D). Crystal structures provide, however, only static conformations of reactants. In contrast, MD simulations unravel the dynamics of the system. In our MD simulations, rotation of γ-phosphate Ras/RasGAP complex brought the O^2G^ atom closer to the P-loop enabling formation of a hydrogen bond with the backbone amide group of Gly13 of the P-loop motif. Formation of this bond was accompanied by elongation of the Gly60-O^2G^ and Gly13-O^3B^ H-bonds in the Ras/RasGAP complex (Fig. 2, S3, Table 1).

Rotation of γ-phosphate into an almost eclipsed conformation should increase the repulsion between phosphorus atoms (Cannon, 1993). Concomitantly, the formation of the new Arg789-O^2G^ H-bond (Fig. 2), is believed to increase the electrophilic nature of the P^G^ atom (Prasad et al., 2013; Wittinghofer & Vetter, 2011). The nucleophilic properties of the attacking water molecule may increase upon rearrangement of the H-bond network after the rotation of γ-phosphate (Jin, et al., 2016; Kamerlin, et al., 2013).

In our MD simulations of the Ras/RasGAP complex, the “catalytic” water molecule provided H-bonds to O^1G^ and Gln61 and accepted a H-bond from the backbone nitrogen of Thr35 (Table 2, Fig. S4). The position of this water molecule is similar to the catalytic water molecule observed in the crystal structure of Ras/RasGAP complex [PDB: 1WQ1] (Scheffzek, et al., 1997), see Fig. S4. The rotation of γ-phosphate (Fig. 2, 3, S2) and its transition into the planar state, by distorting the H-bond with O^1G^, could promote tumbling of the water molecule and formation of a H-bond with the carbonyl oxygen of Thr35, which is just 3.4 Å away (Fig. S4). The H-bond between the backbone nitrogen of Thr35 and the oxygen atom of the catalytic water molecule would then brake, and this oxygen atom would occupy the attack position next to the P^G^ atom. The transition of γ-phosphate into the planar state, however, could not be reproduced by our classical MD simulations. For the Ras-related RhoA GTPase, the stabilization of water molecule in the attack position was recently reported by Blackburn and co-workers who used DFT calculations (Jin, et al., 2016).

The interaction of the amide group of Gly13 both with the O^3B^ atom of the leaving GDP moiety and the O^2G^ atom of γ-phosphate in the Ras/RasGAP complex (Fig. 2, Table 1) might be important for catalysis. Gly13 may initially increase the electrophilicity of the P^G^ atom by interacting with O^3B^ atom (Du & Sprang, 2009) and subsequently stabilize the γ-phosphate upon the further steps of the catalytic transition. The residues that correspond to Gly13 of Ras in diverse P-loop NTPases form H-bonds with the AlF_4_^−^ and AlF_3_ moieties that mimic the planar reaction intermediate (Fig. 1C, D, Tables S3, S4). In our MD simulations, γ-phosphate retained its pyramidal structure. This might be the reason why the H-bond between Gly13 and the O^2G^ atom was not particularly strong (Table 1). In real catalytic reaction, formation of H-bonds between Gly13 and O^2G^, as well as between Arg789 and O^3G^ (Fig. 2A), would drive γ-phosphate towards a more planar, catalytically productive conformation, so that Fig. 3B appears to represent the coordinate of the overall reaction. Further details on the catalysis could be obtained by QM/MM simulations that would incorporate the changes in the geometry of γ-phosphate concomitantly with its rotation, which was out of the scope of this study.

Hence, in the Ras/RasGAP system, rotation of the γ-phosphate group might mechanistically couple weakening of the P^B^-O^3B^-P^G^ bond with an increase in electrophilic properties of the P^G^ atom, transition into a more planar, catalytically productive conformation of γ-phosphate, and polarization of the catalytic water molecule. In this case, GTP hydrolysis would follow one of the currently favored mechanisms, where the bond breakage and the nucleophilic attack proceed in a coupled way (Akola & Jones, 2003; Florian & Warshel, 1998; Plotnikov, et al., 2013).

### Inferring a basal, ancestral mechanism for P-loop NTPases from structure comparison

The mechanism of GTP hydrolysis by H-Ras, inferred from our observations, is consistent with a plethora of data on the functioning of this enzyme, see (Cherfils & Zeghouf, 2013; Kamerlin, et al., 2013; Scheffzek, et al., 1997; Wittinghofer & Vetter, 2011) for reviews. It attributes specific functions to several key elements of the active site that are also found in other classes of P-loop NTPases (Table 2). These are (1) the P-loop motif residues that bind Mg-NTP, stretch the triphosphate chain and partially compensate its negative charges (listed in Table 2 as “P-loop”); some of these residues specifically coordinate γ-phosphate by providing H-bonds that could stabilize the planar pentavalent intermediate (Table 2, “G site”); (2) the activating moiety (such as Arg789 of RasGAP), which provides the positive charge and rotates the γ-phosphate (Table 2, “AG site”), and (3) the “catalytic” residues, such as Gln61 in Ras, which contribute to the binding and coordination of the catalytic water molecule (Table 2, “catalytic”).

Structural analysis (see Fig. 4-6, Table 2, and Supplementary Information) reveals almost unwavering presence of these components across all P-loop NTPases, although the exact nature of the components and the extent of their conservation appear to be different in different classes of P-loop NTPases, see the SI for a detailed description.

The residues of the P-loop that bind Mg-NTP are almost universally conserved, both in sequence and structure (Fig. 6). These residues are present in all known P-loop proteins, with the exception of a few anomalous cases, e.g. the involvement of a His instead of Lys residue in the SAMD9 protein family (Mekhedov et al., 2017), where the His residue can still provide the positive charge normally provided by the conserved Lys residue.

Each class of P-loop NTPases is characterized by its own type of activating “finger” and a distinct mechanism of its insertion. In each case, however, NTP hydrolysis is triggered when the active site with a P-loop-bound NTP molecule comes into contact with another protein domain or subunit. In most cases, a single positively charged group gets inserted between the γ-phosphate and α-phosphate and interacts with both (Fig. 4); then the transition into an eclipsed conformation follows directly from steric considerations: otherwise the O^2A^ and O^3G^ atoms could not simultaneously interact with the activating moiety (Fig. S4). In other instances (Fig. 5), the phosphate chains appear to get distorted by an interaction of the activating moiety solely with the γ phosphate group.

The so-called catalytic/essential residues as well as other γ-phosphate coordinating residues in the G site seem to be auxiliary to the activation of NTP hydrolysis, as they vary between proteins within each class and can be even absent in certain functional NTPases (Table 2). For instance, certain AAA+ proteins provide no additional support for the γ-phosphate group, see Table 2 and (Wendler, et al., 2012) for review.

The strict conservation of the key components within P-loop NTPases (Table 2) indicates a common basal mechanism of NTP hydrolysis. Even apparent exceptions, such as the ABC transporters, strengthen the case for such a basal mechanism by identifying the genuinely compulsory elements. ABC transporters show that the role of the activating moiety cannot be reduced solely to its electrostatic contribution, as is widely believed. In ABC transporters, the electrostatic impact of two protein backbone amide groups can hardly match that of an Arg finger, and yet the LSGGQ motif can trigger ATP hydrolysis via an interaction with γ-phosphate.

### Roles of the residue in the Gly13 position of Ras GTPases

Here, based on structure analysis (Fig. 1C, D, Table S3, S4) and MD simulations (Fig. 2, S3, Table 1), we argue that Gly13 of Ras, as well as the corresponding residues in other P-loop NTPases, could, in addition to their interaction with the bridging O^3B^ atom, also stabilize the O^2G^ atom of γ-phosphate after its rotation into the catalytically productive eclipsed conformation and promote the planar geometry of γ-phosphate. The potential importance of the H-bond between this backbone nitrogen and O^2G^ for catalysis is justified by the presence of an analogous bond in the structures of diverse P-loop NTPases that were crystallized with transition state analogs or non-hydrolizable NTP analogs (Table S3, S4, Fig. 1C,D). This H-bond is particularly frequently seen in the structures that contain NDP-AlF_4_^−^ complex, the closest mimic of the pentavalent intermediate (Table S3). The position of the backbone amide group of the counterparts of Gly13 in the vicinity of the O^2G^ atom is structurally conserved across P-loop NTPases (Fig. 6A), being determined by the highly conserved H-bond with the bridging O^3B^ oxygen (Fig. 1A, 2, S3, Table 1). Although only the backbone of Gly13 (or its counterparts) is involved in the interaction with O^2G^ of GTP, this position is consistently occupied by either Gly or other small (Ala or Ser) residue in most classes of P-loop NTPases (Table 2, S3, S4, Fig. 6B). The presence of small residues should enable the flexibility of the backbone. Conservation of the small residues in the Gly13 position in most P-loop NTPases (Fig. 6B) points to their interaction with an invariant part of the catalytic machinery, which appears to be the O^3B^ and O^2G^ atoms of the phosphate chain. Therefore, we believe that the flexibility of the backbone at Gly13 position might be important for the ability of the backbone amide to interact with O^3B^ and/or O^2G^ atoms also in other families of P-loop NTPases.

Among crystal structures with a H-bond between counterparts of Gly13 and AlF_4_^−^, larger residues are seen in the Gly13 position only in a few NTPase families, all of them within TRAFAC class (Table S3). In cation-dependent TRAFAC NTPases, the Gly13 position is taken by an Asn or Asp residue that coordinates the activating K^+^ ion, see Table 2, Fig. 4B, 5A, (Anand, et al., 2010; Ash, et al., 2012; Mulkidjanian et al., 2012b; Scrima & Wittinghofer, 2006; Wuichet & Sogaard-Andersen, 2015) and (Shalaeva et al., submitted); the backbone nitrogen atoms of these residues make H-bonds with the AlF_4_^−^ moieties in the respective structures (Table S3). It appears that the activating K^+^ or Na^+^ ion not only interacts with the O^3G^ atom of γ-phosphate (see Fig. 4B, 5A), but also, indirectly, “communicates” with the O^2G^ atom via the backbone nitrogen atom of Asn/Asp in the Gly13 position; this interaction may be functionally relevant.

We have previously presented evidence suggesting that activating Arg resudues were independently recruited, instead of K^+^ ions, in diverse lineages of TRAFAC NTPases (Dibrova et al., 2015; Mulkidjanian, et al., 2012b), which implies that the common ancestor of TRAFAC NTPases was a K^+^-dependent enzyme with an Asp/Asn in the Gly13 position. Accordingly, residues congruent to Asp/Asn could have remained in the Gly13 position even in some TRAFAC NTPases that have lost the K^+^-dependence. Indeed, in the small GTPase Arf-like 3 protein (Arl3), an Asn in the same position is turned away from the AG site, which accommodates an Arg finger, and makes an additional H-bond with the activating protein (Veltel et al., 2008). The G_α_ subunits of heterotrimeric G-proteins have, in the respective position, a Glu residue that, after GTP hydrolysis, interacts with the liberated Arg finger to form a salt bridge that locks the GDP molecule in the binding site (Wall et al., 1995).

These structural traits explain the obscure fact that the Gly13Asp mutation in Ras GTPases is by far the most oncogenic (Bos, et al., 1987; Prior, et al., 2012; Wey, et al., 2013). The carboxyl group of Asp13, by occupying the AG site, should interact with the Arg789 finger (perhaps, even making a salt bridge with it), hinder hydrolysis, and thereby prevent cancellation of the oncogenic signal. The far less frequent oncogenicity of some other mutations of Gly13, such as Gly13Cys or Gly13Val (Bos, et al., 1987; Prior, et al., 2012; Wey, et al., 2013), could be due to an insufficient flexibility of the backbone in the absence of Gly13.

### Relation to the previous studies of P-loop NTPases

The rotation of the γ-phosphate group and formation of an additional H-bond with Gly13 in H-Ras/RasGAP complex, which are seen not only in our MD simulations (Fig. 2), but also in many crystal structures that contain transition state analogs or non-hydrolizable NTP analogs (Fig. 1C, D, Table S3, S4), were so far overlooked in simulation studies. We suspect that the reason was the insufficient simulation time. Earlier MD simulations were done on a picosecond time scale (Ma & Karplus, 1997; Resat, et al., 2001). The more recent long-lasting MD simulations of diverse Ras proteins addressed questions other than the catalytic mechanisms (Lu, Jang, Muratcioglu, et al., 2016; Prakash & Gorfe, 2014). The catalysis by H-Ras/RasGAP, in recent studies, has been mostly scrutinized by sophisticated QM/MM simulations, which are also performed on the time scale of only picoseconds (Jin, et al., 2016; Khrenova, et al., 2015; Prakash & Gorfe, 2014; Rudack, et al., 2012). Our 3×10 ns MD simulations appear to be among the longest ever applied to both H-Ras and Ras/RasGAP complex with a goal of clarifying the catalytic mechanism.

For the Ras GTPase and heterotrimeric G proteins, Gerwert and coworkers proposed that the Arg finger, by rotating the α-phosphate group relatively to the β- and γ-phosphates, brings the phosphate chain of small G-proteins into a fully eclipsed conformation (Gerwert, et al., 2017; Mann, et al., 2016; Rudack, et al., 2012). Their suggestion was based on the results of QM/MM simulations of methyltriphosphate in water (Rudack, et al., 2012), where the rotation of the α-phosphate was not restricted. We, while estimating the energy of K^+^ binding to the AG site of a Mg-ATP complex in water (Shalaeva et al., submitted), have also observed that binding of the second cation in the AG site caused a major rotation of the α-phosphate that, together with a minor rotation of γ-phosphate, yielded a fully eclipsed conformation of the phosphate chain. In P-loop-bound NTP molecules, however, rotation of α-phosphate is prevented by interactions of the whole NTP molecule with the enzyme. Both in the Ras and Ras/RasGAP crystal structures, α-phosphate forms a stable H-bond with the backbone amide group of Ala18, these bonds persisted in all our MD simulations (Fig. 2, S3, Table 1). The shape of the P-loop and the location of α-phosphate are extremely well conserved across all P-loop NTPases (Fig. 6, Table 2), indicating that a similar bond exists in other P-loop NTPases. Still, even the breakage of this H-bond would not permit rotation of α-phosphate because the moieties from the both sides of α-phosphate (the nucleobase-ribose and Mg-bound β- and γ-phosphates, respectively) are tied by multiple interactions to the protein (Fig. 1A, 3, 6). Therefore, the rotation around the bridging bonds of P^A^ is hindered; the neighboring moieties cannot rotate within the active site, especially when the activating protein/domain (e.g. RasGAP) is bound.

Our MD simulations showed that the near-eclipsed conformation of the phosphate chain was achieved mostly via the rotation of γ-phosphate (Fig. 2, 3). In contrast to α-phosphate, γ-phosphate, being the end group, can rotate even within the complex between the P-loop domain and its activator (Fig. 1B-D). The rotation of γ-phosphate results not only in an almost eclipsed conformation, but also in reshuffling of the H-bond network around γ-phosphate and in its more planar state, which would drive the catalytic reaction.

Our comparative structure analysis supports the data of Ogura and coworkers on the functional importance of the Arg residues in the active sites of the AAA and AAA+ families (Ogura, et al., 2004). Furthermore, our structure analysis indicates that electrophilic activating moieties are involved in all classes of P-loop NTPases (Fig. 4, 5).

Our results also agree with the suggestion of Warshel and coworkers that the common feature in P-loop NTPases is the electrostatic stabilization of the transition state (Prasad, et al., 2013). The H-bond between Gly13 and the O^3G^ should contribute to such stabilization upon the catalytic transition.

In contrast, the suggestion of Kiani and Fischer on the general importance of catalytic Glu residues in polarizing the attacking water molecule (Kiani & Fischer, 2016) could not be confirmed: Glu residues, as holders of catalytic water molecules, are common only for the ATPases of the ASCE division with their position varying between classes, see Table 2 and SI. The poor conservation of “catalytic” residues (Table 2) strongly supports the suggestion of Warschel and coworkers (Schweins, et al., 1994) that polarization/deprotonation of the attacking water molecule is mediated by γ-phosphate itself, whereas the function of the “catalytic” residue might be just to hold the water molecule in the proper position.

Our data, in essence, agree with the suggestion of Blackburn and coworkers that the activating Arg residue, in Ras-related NTPases, causes the reshuffling of H-bonding around the catalytic water molecule and forces it into the attacking position. In our MD simulation of the Ras/RasGAP complex, the water molecule close to the attack position was coordinated by Thr35 and Gln61 (Fig. S4); corresponding residues were suggested by Blackburn and coworkers to position the attacking water molecule in the Ras-related RhoA GTPase (Jin, Molt, et al., 2017; Jin, et al., 2016; Jin, Richards, et al., 2017).

### Energetics of P-loop NTPases

The function of P-loop NTPases, as discussed above, is not to catalyze the NTP hydrolysis *per se*, but to do so only in response to an interaction with an activating protein. However, there is always a possibility that a protruding Arg/Lys residue of some unrelated protein or an occasionally bound K^+^ ion could cause non-specific activation of a P-loop NTPase. As discussed in (Shalaeva et al., submitted), the danger of spurious activation is particularly high in case of K^+^(Na^+^)-activated GTPases because the cation-binding K-loop is part of the P-loop domain. Hence, activation should be specific, and take place only upon the interaction with a proper partner.

Our MD simulations showed that insertion of an activating cationic moiety in the AG site is coupled with a notable energy barrier. Fig. S6 shows that free energy of about 20-25 kJ/mol appears to be needed to rotate the γ-phosphate towards the eclipsed conformation, which is sterically unfavorable (Cannon, 1993). Therefore, NTP hydrolysis by a dedicated activating protein requires free energy to impel the K^+^ ion or an Arg/Lys residue into the AG site and to twist the γ-phosphate. The calculated free energy value is comparable with the decrease of the activation barrier in the presence of RasGAP. Indeed, the interaction with RasGAP accelerates GTP hydrolysis by five orders of magnitude as compared to Ras alone, which corresponds to decreasing the activation barrier by ca. 27-30 kJ/mol (Glennon et al., 2000; Kotting & Gerwert, 2004). Hence, this acceleration appears to be mostly achieved owing to the rotation of the γ-phosphate by Arg789 (20-25 kJ/mol, Fig. S6), which is in agreement with the experimental data on the insertion of Arg789 as the rate-limiting event of the catalytic reaction (Kotting & Gerwert, 2004).

The specific problem of P-loop NTPases is the source of free energy for activation. In enzyme catalysis, free energy for decreasing the activation barrier - by destabilizing the substrate and/or stabilizing the transition state - is typically gained from substrate binding (Fersht, 1985; Jencks, 1987). In the case of P-loop NTPases, where the substrate binding step is separated from the ultimate catalytic step, the energy of substrate binding to the P-loop is used to yield a catalytically prone conformation of the bound substrate with eclipsed β- and γ-phosphates (Delbaere, et al., 2004; Matte, et al., 1998). While increasing the rate of hydrolysis by several orders of magnitude as compared to that in water, the free energy of NTP binding to the P-loop is used without achieving physiologically relevant hydrolysis rates (Kotting & Gerwert, 2004; Shutes & Der, 2006). For fast hydrolysis, an additional source of free energy is needed (27-30 kJ/mol in the case of Ras).

The need for an additional input of energy has been widely discussed in relation to the oligomeric, ring-forming P-loop NTPases, such as AAA+ ATPases (Hattendorf & Lindquist, 2002; Lupas & Martin, 2002) and rotary ATPases/helicases (Boyer, 1997; Senior et al., 2002) (Table 2, Fig. 4 D-F). In these enzymes, the energy for inserting the arginine finger into the ATP-containing catalytic site is provided by the binding of another nucleotide molecule to the other monomer. The free energy of nucleotide binding in the other site drives the conformational change that is transmitted to the Arg finger. Hence, insertion of an activating finger into the AG site (Fig. 4D-F) proceeds not owing to the substrate binding in this site, but at the expense of the binding energy that is gained somewhere else in the enzyme complex and delivered via conformational coupling. However, many P-loop NTPases, and specifically, small regulatory GTPases, are monomeric and only interact either with their activating proteins or with each other upon dimerization (Table 2). By exclusion, only the free energy of their binding to the activating partner is available to serve as the driving force for impelling the activating positive charge, either a K^+^ ion or an Arg/Lys residue, into the AG site. Evidence is available that the catalytic activity of some P-loop NTPases is indeed controlled by the strength of their binding to the activating proteins (Cherfils & Zeghouf, 2013; Codina & Birnbaumer, 1994). How exactly the energy of protein-protein binding is used to push the activating moiety into the AG site is yet to be established and is likely to be specific for each P-loop family. It also remains to be established what is the function of the P-loop fold in mediating the energy funneling. One consequence of such a mechanism is that by changing the energy of interaction between the two proteins, it could be possible to change the rate of the catalytic reaction. There are numerous ways to affect the interaction between two proteins - e.g. by invoking more protein partners, which is a typical way to regulate P-loop NTPases (Table 2). Because of this mechanism, NTPases are easily tunable and versatile.

### Shared catalytic mechanism of P-loop NTPases

We propose here a two-step common mechanism for the whole superfamily of P-loop NTPases.

1. Upon NTP binding to the P-loop, the energy of binding is used to bring the NTP molecule into a catalytically prone conformation with eclipsed β- and γ-phosphates via coordination of the Mg^2+^ ion and phosphate groups by multiple amino acids that are strictly conserved among all P-loop NTPases (Fig. 1, 4-6, Table 2).
2. NTP hydrolysis is activated by a specific interaction of the P-loop NTPase with another protein or a separate domain of the same protein or a RNA/DNA molecule. This interaction leads to the insertion of a positively charged moiety next to the phosphate chain. In most cases, this moiety is an Arg/Lys residue or a monovalent cation. The activating moiety either bridges O^2A^ and O^3G^ (Fig. 4A-G) or interacts solely with O^3G^ (Fig. 5A-F). Either way, this interaction, via rotation/pulling of the γ-phosphate group, yields a near-eclipsed conformation of the whole phosphate chain, which causes a further weakening of the P^B^-O^3B^-P^G^ bond and a more planar state of γ-phosphate. The reshuffling of the H-bond network around γ-phosphate increases the electrophilic nature of the P^G^ atom and brings the catalytic water molecule into the attacking position where it is typically supported by the “catalytic”/“essential” residues. The cleavage of the P^B^-O^3B^-P^G^ bond proceeds by a mechanism where weakening of this bond and the nucleophilic attack are coupled via the motion of γ-phosphate.

The proposed universal mechanism of NTP hydrolysis by P-loop NTPases, the most widespread catalytic mechanism in living nature, appears to be pretty simple and robust. In a way, removal of γ-phosphate from an NTP molecule resembles plucking an apple from a tree – the “fingers” are used to forcibly rotate the γ-phosphate and then pull it off.

The free energy for twisting/pulling the γ-phosphate (ca. 20-25 kJ/mol in case of Ras/RasGAP, Fig. S6) could be provided by the interaction between the P-loop protein domain and the activating protein domain. Hence, the rate of hydrolysis by P-loop NTPases could be fine-tuned by affecting protein-protein interactions. We suggest that this easily tunable and versatile mechanism of using the free energy of protein-protein interaction for catalysis provides unlimited possibilities to control NTP hydrolysis, which might explain why P-loop ATPases and GTPases, despite their limited catalytic repertoire, represent the most common protein fold.

### Evolutionary considerations

The abundance and diversity of P-loop NTPases has long obscured the evolutionary origins of this vast superfamily. It is noteworthy that the major classes of P-loop NTPases can be traced back to the Last Universal Cellular Ancestor (LUCA), the common ancestor of bacteria and archaea (Baldauf et al., 1996; Gogarten et al., 1989; Iwabe et al., 1989; Leipe, et al., 2003; Leipe, et al., 2002; Wuichet & Sogaard-Andersen, 2015). The current view on the origin of P-loop NTPases is that they have evolved from an NTP-binding protein which, like the P-loop motif itself, was incapable of fast NTP hydrolysis (Alva, et al., 2015). The common catalytic mechanism of P-loop NTPases, proposed in this work, offers a simple explanation for the conservation of the key residues of the P-loop motif. It posits that the residues that initially participated in binding the NTP triphosphate chain continued doing so over the entire evolutionary history of this superfamily. In contrast, the activating moieties were under no such constraint: in the absence of an activator, there was very little or no NTP hydrolysis. Accordingly, the activating moieties rapidly diverged to produce the current diversity of the P-loop classes and families. As detailed in Table 2, each of those classes has its own, specific mechanism of providing one or more positive charges to the NTPase active site; the TRAFAC class that has several such mechanisms.

As argued above and in (Shalaeva et al., submitted), in the NTPases of the TRAFAC class, the presence of K-loop alone is not sufficient for (fast) NTP hydrolysis. Therefore, even if an ancestral P-loop NTPase occasionally attained some ancestral version of the K-loop, the low intrinsic affinity of the AG site to K^+^ should have prevented it from functioning as a catalytic cofactor. The affinity to K^+^ ions could, however, increase upon binding of the ancestral P-loop protein to its functional partner, e.g. the ribosome. The K-loop could then be fixed in a conformation that would force the K^+^ ion to enter the AG site. As a result, (i) the energy of the binding of a P-loop protein to its partner could be utilized to drive the NTP hydrolysis and (ii) the NTP hydrolysis could be timed to the binding event. Furthermore, the rate of catalysis could be modulated by changing the specificity and/or affinity of the binding. The potency of this mechanism for regulation of biological processes can hardly be overestimated.

The eventual recruitment of positively charged Arg/Lys fingers made the enzymes independent of K^+^ ions and therefore enabled an even better control over the catalytic reaction.

The functional replacements of K^+^ ion by Lys/Arg fingers are so widespread because the underlying adaptive changes do not appear to be particularly demanding. Any Lys/Arg residue with its long and flexible side chain reaching the triphosphate moiety would do the job, provided that it is appropriately pushed by the respective binding interaction. As a result, activation by Arg or Lys fingers is used, in different modifications, in many enzymes and makes the basis of biological regulation and control over the enzyme activity via inserting or removing the charged residue, e.g. as a result of protein-protein interaction, protein dimerization or otherwise induced conformational changes.

The insertion of Arg/Lys fingers, similarly to K^+^ ions, could lead to activation only if driven by a thermodynamically favorable event, usually an interaction with another protein or domain (e.g. upon dimerization), or an RNA/DNA molecule. Further elaboration of the mechanism in case of regulatory GTPases (TRAFAC) was coupled with the involvement of further regulatory proteins, each of which modulated the binding between the P-loop GTPase and its activator and, in this way, controlled the catalytic reaction.

Hence, our analysis suggests that separation of the ancestral P-loop fold into two divisions and 8 classes has occurred even before the LUCA. The K^+^-activated P-loop NTPases, specifically of the TRAFAC class, could have emerged only if potassium ions prevailed over sodium ions in the habitats of the first cells; otherwise, Na^+^ ions would always bind instead of K^+^ ions. Thus, our results support the earlier suggestion that the first cells have been shaped in K^+^ rich environments, such as the primordial anoxic geothermal fields (Mulkidjanian et al., 2012a; Mulkidjanian, et al., 2012b).

## Methods

The conformational mobility of GTP in complex with Ras and Ras/RasGAP was studied by molecular dynamics (MD) simulations. Each protein complex was placed in a cubic cell filled with TIP3P water with standard periodic boundary conditions. Minimal distance between any atom of the protein and the periodic cell wall was set at 12 Å. A mixture of Na^+^ and Cl^−^ ions were randomly placed in both systems to bring the total charge to neutrality. For simulations, we used the CGenFF v.2b8 force field for the ATP and GTP and CHARMM36 force field for the protein (Vanommeslaeghe, et al., 2010). For the Mg^2+^ ion we used parameters designed by Calahan et al. (Callahan et al., 2010). Equilibration of each system was achieved by a series of simulations with restraints on the conformation of the protein and free MD simulations with the duration of 10 ns. Productive runs were performed with constant temperature and pressure (NPT ensemble) with the temperature of 303.15 K, controlled by Nose-Hoover thermostat, and the pressure of 1 atm, controlled by Parrinello-Rahman barostat. Long-range electrostatic interactions were computed using particle mesh Ewald method with Verlet cutoff-scheme (Verlet, 1967) and a 12 Å distance cutoff for direct electrostatic and van der Waals interactions. Switching function was set to 10 Å to gradually reduce van der Waals potentials, reaching 0 at the cutoff distance. The geometry of bonds between hydrogens and heavy atoms was constrained to the lengths and angles defined by the force field using LINCS (LINear Constraint Solver) algorithm (Hess et al., 1997). For each of the two systems, three independent unconstrained simulations of 10 ns each were performed (productive runs). During simulations, conformations of the GTP molecule were saved every 5 ps, and conformations of the entire system were saved every 100 ps. Productive runs were used to extract characteristic frames to represent the geometry of the GTP binging site with visual molecular dynamics (VMD, (Humphrey et al., 1996)) and to calculate statistics of structure movement using MATLAB R2017a (“MATLAB and Statistics Toolbox Release 2017a,” 2017).

Comparative structural analyses that included structure superposition and distance determinations was performed in PyMol v 1.7.2.1 (2010), which was also used to draw Fig. 1-6.

## Acknowledgements

Very useful discussions with Drs. A.V. Golovin, A. Gorfe, Y. Kalaidzidis, J. Klare, E.V. Koonin, V.P. Skulachev and H.-J. Steinhoff are greatly appreciated. We are thankful to Dr. D. Dibrova and A. Mulkidzhanyan for their help during the launching phase of this project. This study was supported by the Deutsche Forschungsgemeinschaft, Federal Ministry of Education and Research of Germany (A.Y.M.), the German Academic Exchange Service (D.N.S.) a grant from the Russian Science Foundation (14-50-00029, D.N. S., D.A.C), and the Lomonosov Moscow State University (Supercomputer Facility, D.A.C). M.Y.G. is supported by the Intramural Research Program of the NIH at the National Library of Medicine.

## Supporting Information

### Supplementary Tables

**Table S1.**
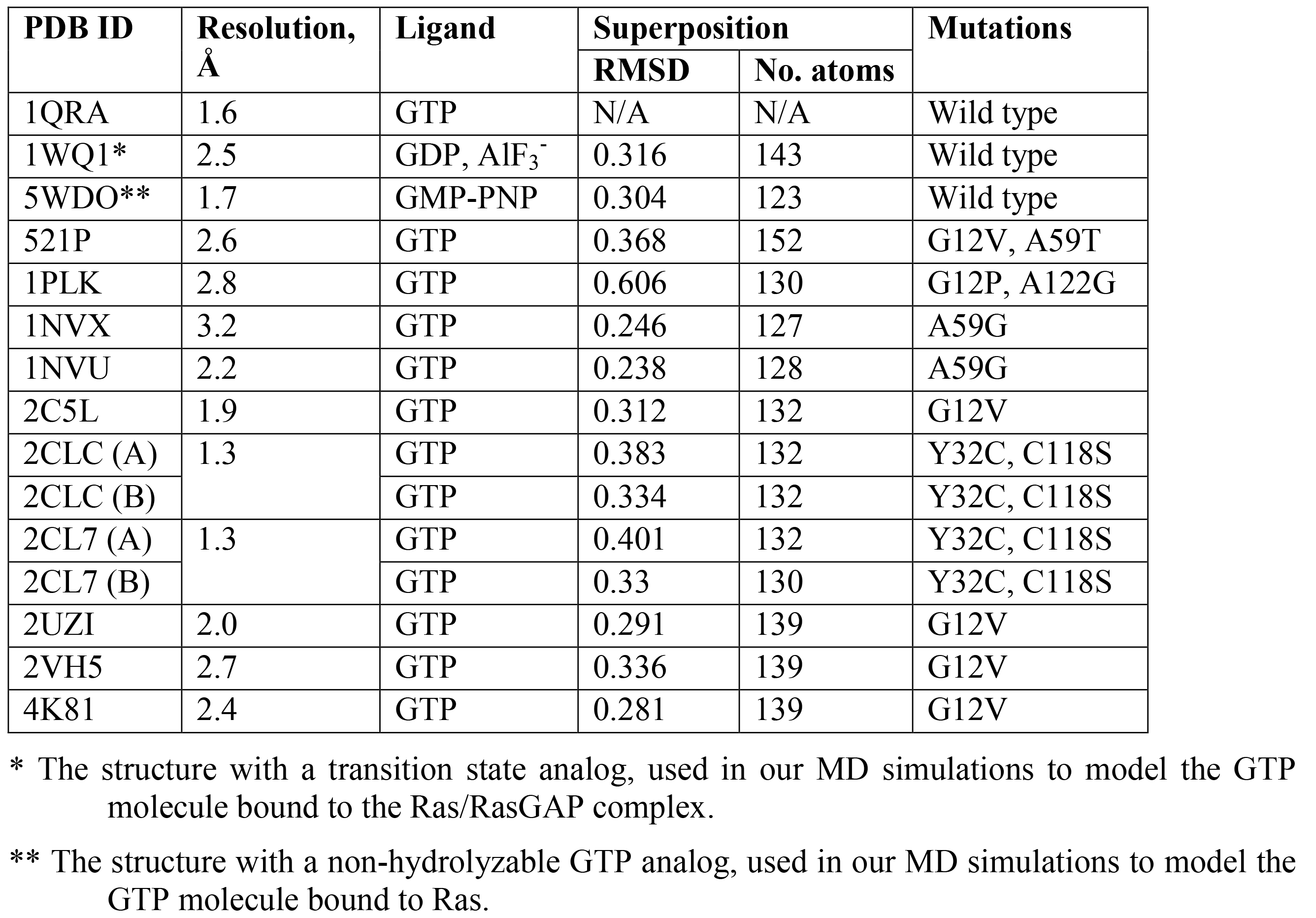
Crystal structures of Ras GTPase in complex with Mg-GTP.

| PDB ID | Resolution,<br>Å | Ligand | Superposition |  | Mutations |
| --- | --- | --- | --- | --- | --- |
|  |  |  | RMSD | No. atoms |  |
| 1QRA | 1.6 | GTP | N/A | N/A | Wild type |
| 1WQ1* | 2.5 | GDP, AlF<sub>3</sub><sup>-</sup> | 0.316 | 143 | Wild type |
| 5WDO** | 1.7 | GMP-PNP | 0.304 | 123 | Wild type |
| 521P | 2.6 | GTP | 0.368 | 152 | G12V, A59T |
| 1PLK | 2.8 | GTP | 0.606 | 130 | G12P, A122G |
| 1NVX | 3.2 | GTP | 0.246 | 127 | A59G |
| 1NVU | 2.2 | GTP | 0.238 | 128 | A59G |
| 2C5L | 1.9 | GTP | 0.312 | 132 | G12V |
| 2CLC (A) | 1.3 | GTP | 0.383 | 132 | Y32C, C118S |
| 2CLC (B) |  | GTP | 0.334 | 132 | Y32C, C118S |
| 2CL7 (A) | 1.3 | GTP | 0.401 | 132 | Y32C, C118S |
| 2CL7 (B) |  | GTP | 0.33 | 130 | Y32C, C118S |
| 2UZI | 2.0 | GTP | 0.291 | 139 | G12V |
| 2VH5 | 2.7 | GTP | 0.336 | 139 | G12V |
| 4K81 | 2.4 | GTP | 0.281 | 139 | G12V |

\* The structure with a transition state analog, used in our MD simulations to model the GTP molecule bound to the Ras/RasGAP complex.
\*\* The structure with a non-hydrolyzable GTP analog, used in our MD simulations to model the GTP molecule bound to Ras.

**Table S2.** Dihedral angle values in the phosphate chains of Mg-GTP in H-Ras and Ras/RasGAP.

| System | Structure | $\Psi^{\alpha-\beta}$ | $\Psi^{\beta-\gamma}$ | $\Psi^{\alpha-\gamma}$ |
| --- | --- | --- | --- | --- |
| Mg-GTP-Ras | PDB:<br>1QRA | 49° | 6° | 49° |
| Mg-GTP-Ras | MD<br>simulation | 59±6° | 17±8° | 69±10° |
| Mg-GDP-AlF3-Ras/RasGAP | PDB:<br>1WQ1 | 46° | -7° | 49° |
| Mg-GTP-Ras/RasGAP | MD<br>simulation | 46±6° | -6±11° | 33±12° |

**Table S3.**
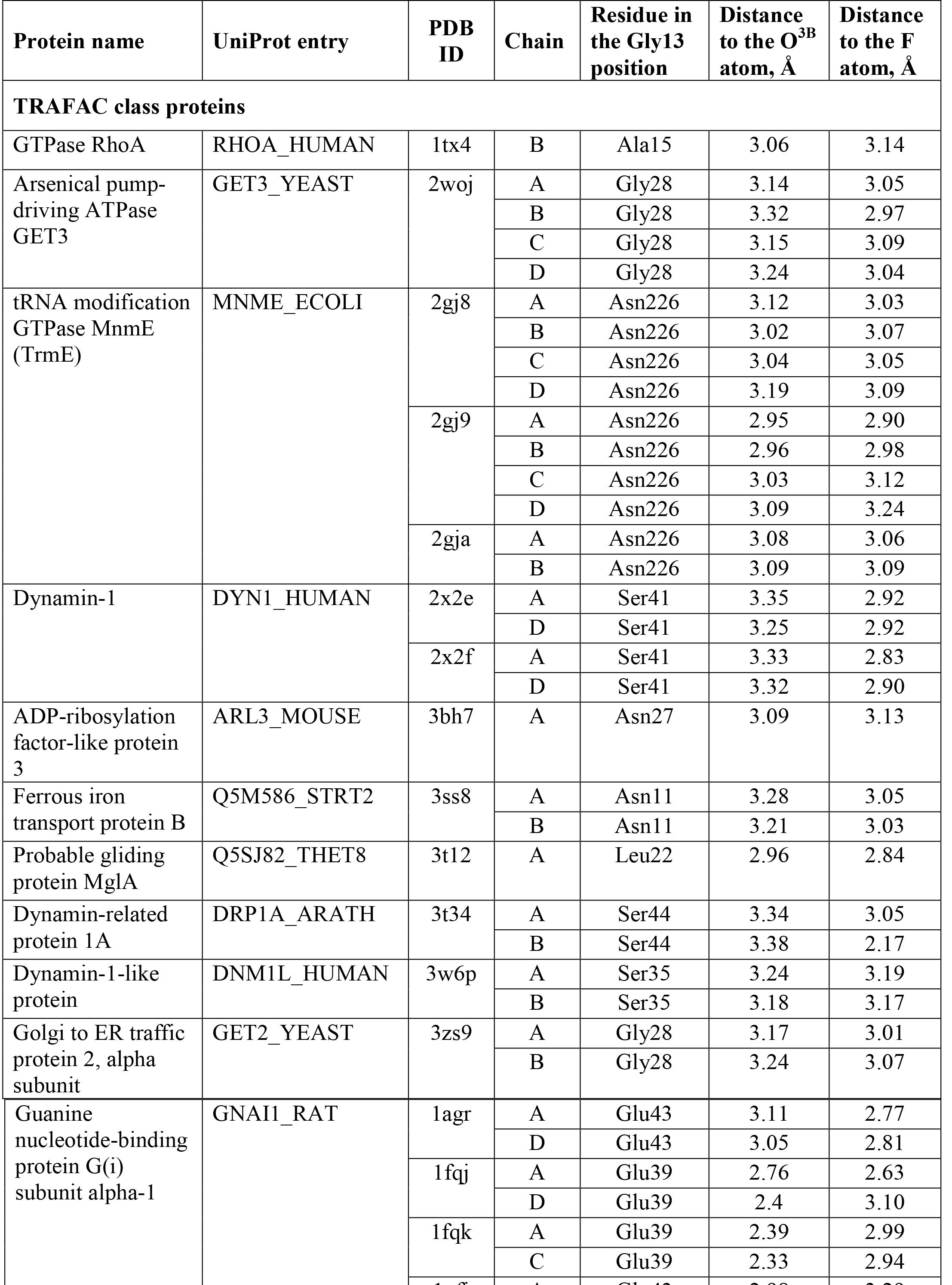
Interactions of the residues corresponding to Gly13 of Ras with AlF_4_^−^ moieties used in transition state analog complexes

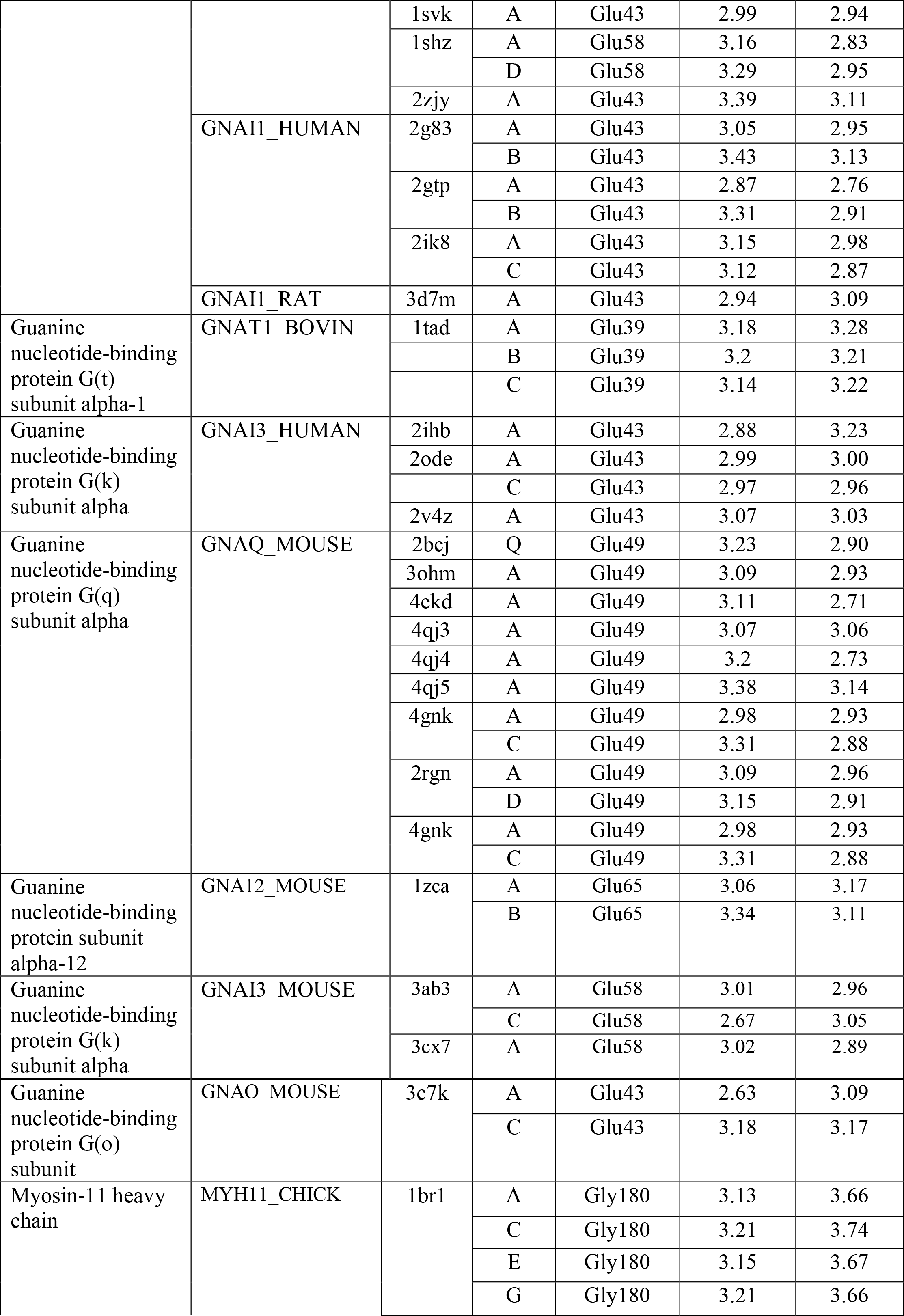

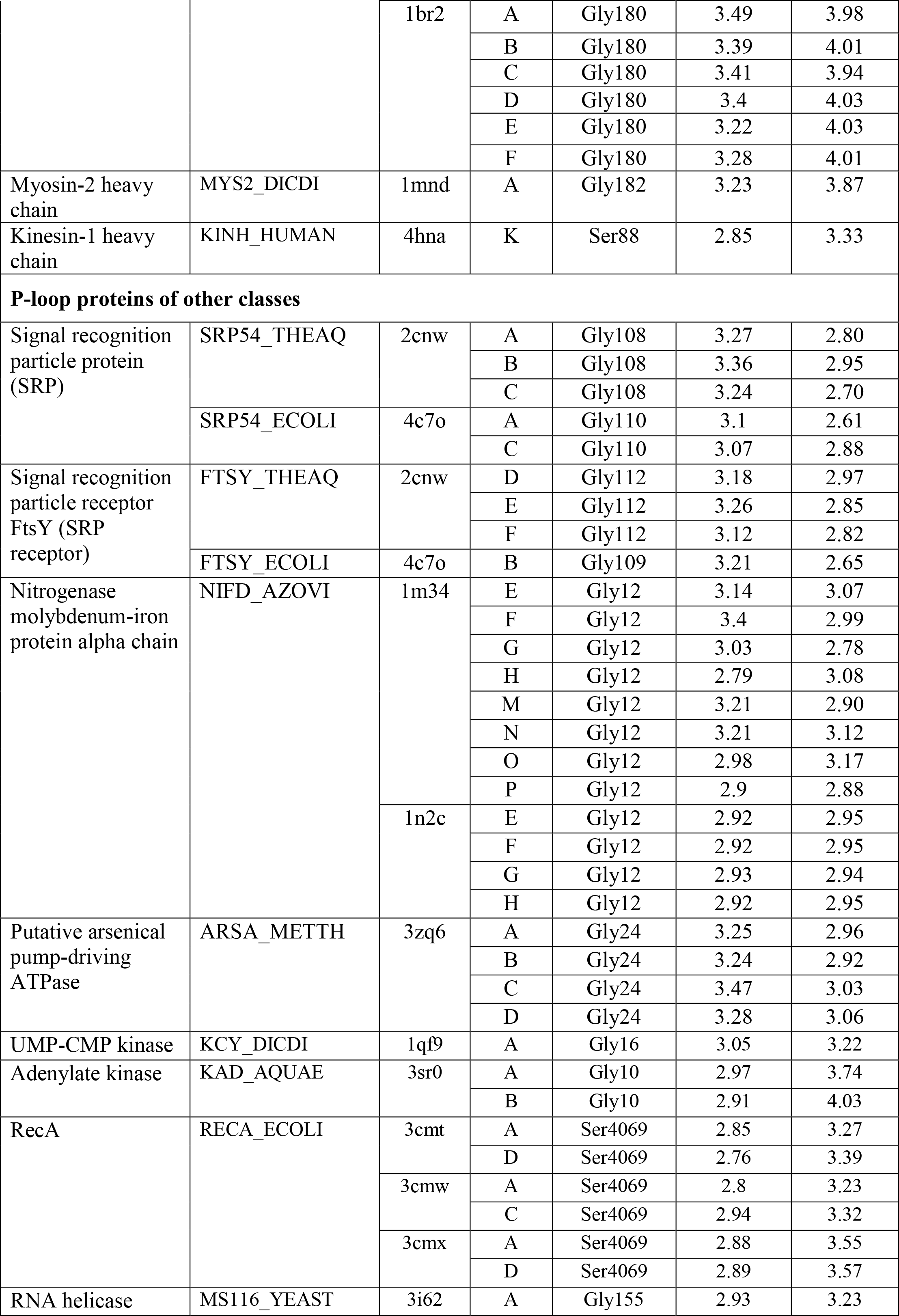

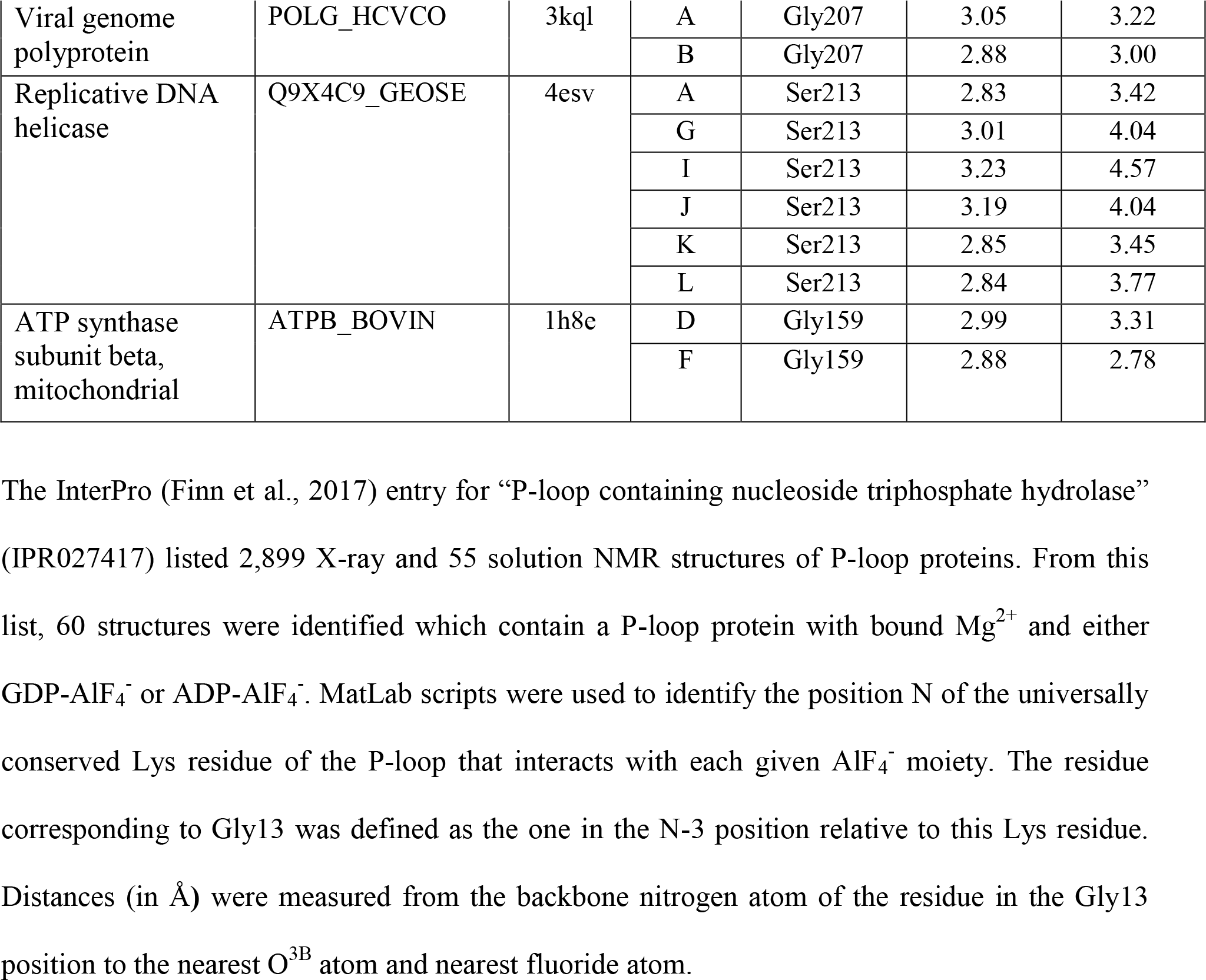

**Table S4.**
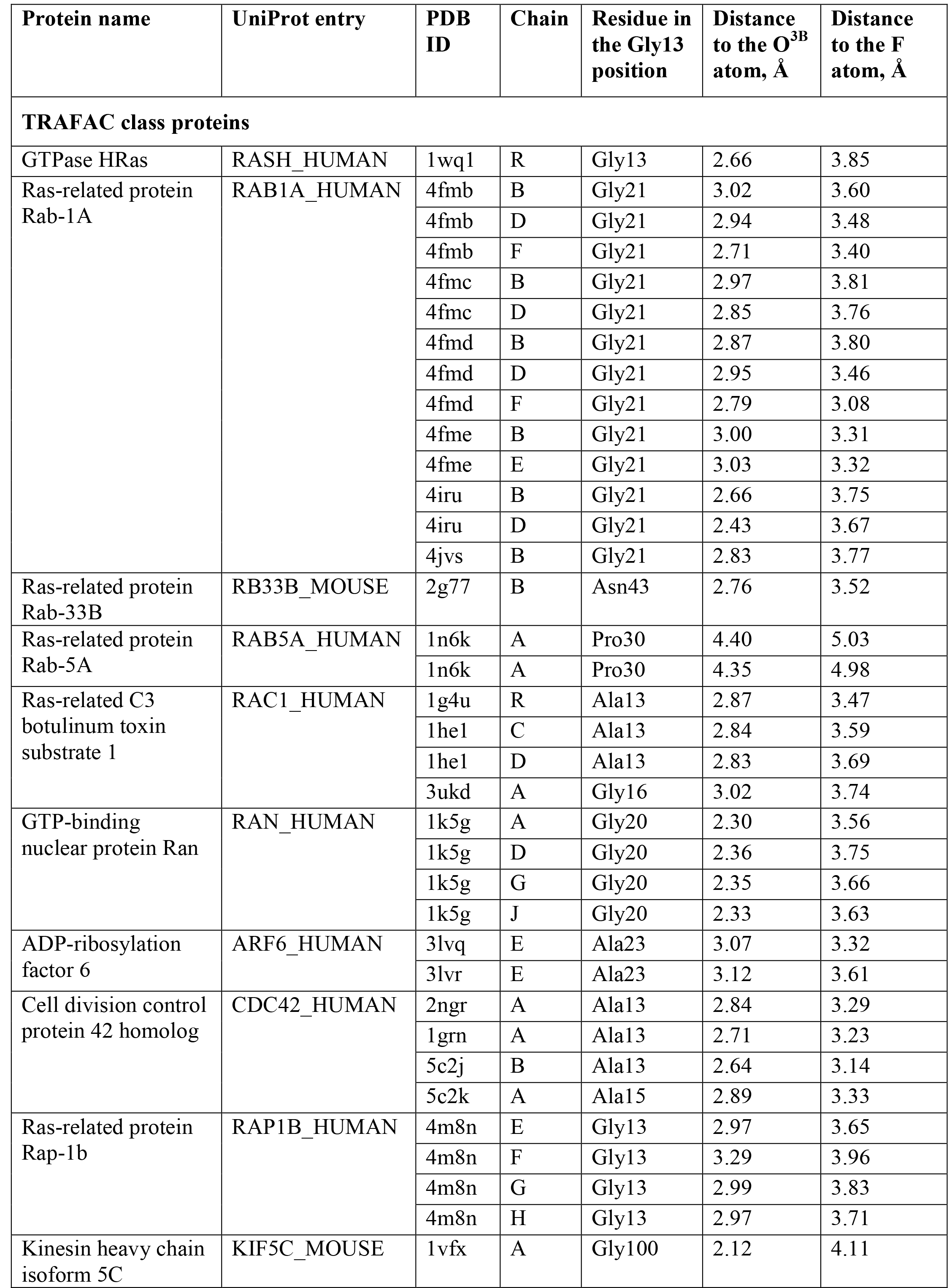
Interactions of the residues corresponding to Gly13 of Ras with AlF_3_ moieties used in transition state analog complexes

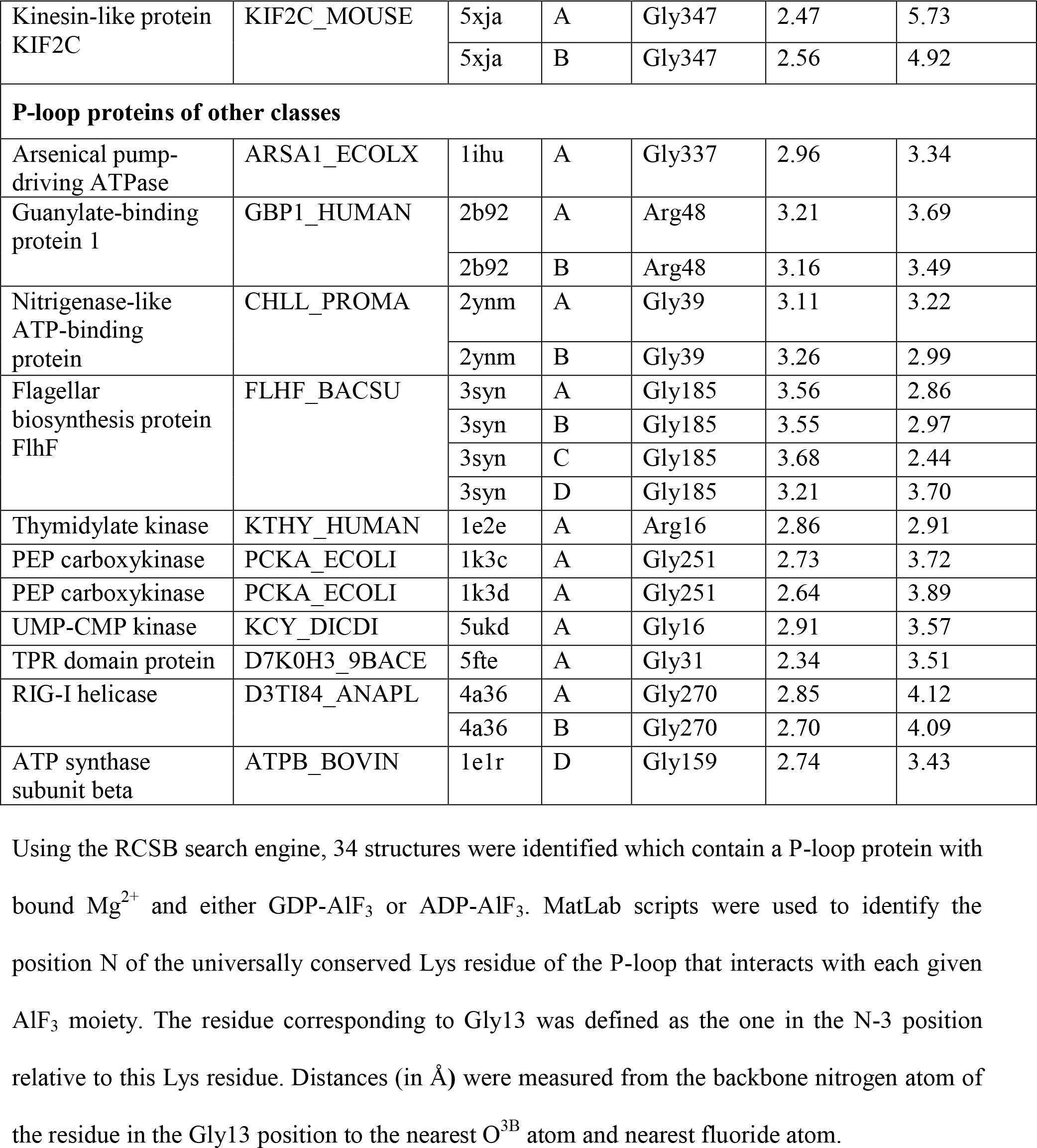

### Supplementary Figures

**Figure S1.**
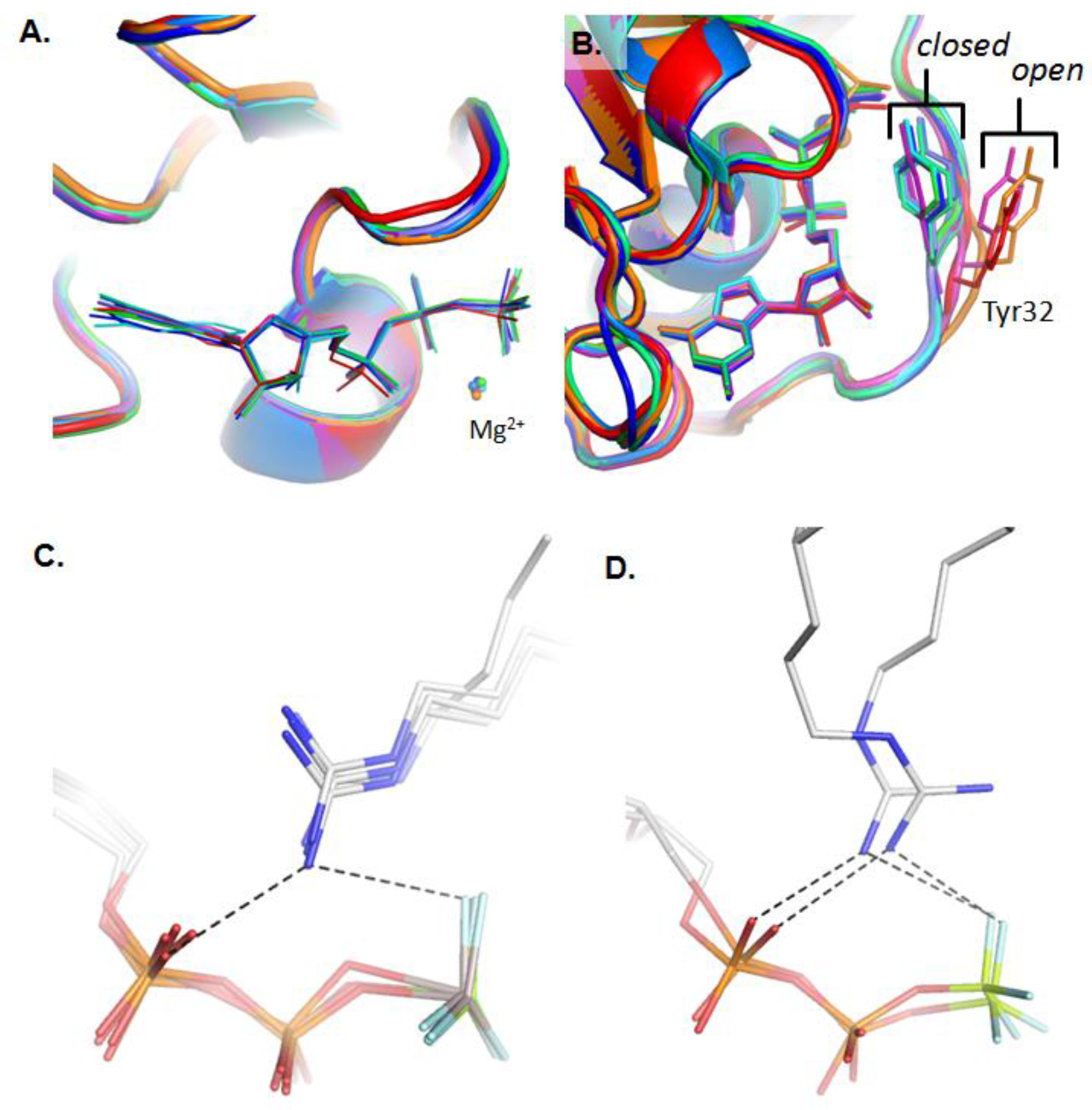
Crystal structures of Ras and Ras-like proteins in complex with GTP and its analogs. **A.** Conformations of GTP molecules in the active site of Ras. The superimposed PDB entries are 1QRA, 521P, 1NVX, 1NVU, 2C5L, 2UZI, 2VH5, and 4K81. GTP is shown as sticks, Mg^2+^ ions shown as spheres. **B.** Open and closed conformations of Tyr32 in Switch I. Structures with the open conformation are 1QRA, 521P, and 1WQ1; structures with closed conformation are 1NVX, 1NVU, 2C5L, 2UZI, 2VH5, 4K81, and 5WDO. GTP and Tyr32 are shown as sticks, Mg^2+^ ions shown as spheres. **C.** Interaction of Arg fingers from GAPs with GDP-AlF_3_ and GDP-MgF_3_ in Ras-like protein complexes: RhoA/ArhGAP35 [PDB 5IRC], RhoA/ArhGAP20[PDB 3MSX], Cdc42/RacGAP1 [PDB 5C2J], and Rac1/exoS [PDB 1HE1]. Black dashes indicate distances in the range of 2.6-3.3 Å. **D.** Interaction of Arg fingers from GAPs with GDP-BeF_3_ in Rab GTPase protein complexes: Rab-1B/RabGAP (TBC1D20) [PDB 4HLQ] and Rab-1B/LepB [PDB 4I1O]. Black dashes indicate distances in the range of 2.6-3.3 Å.

**Figure S2.**
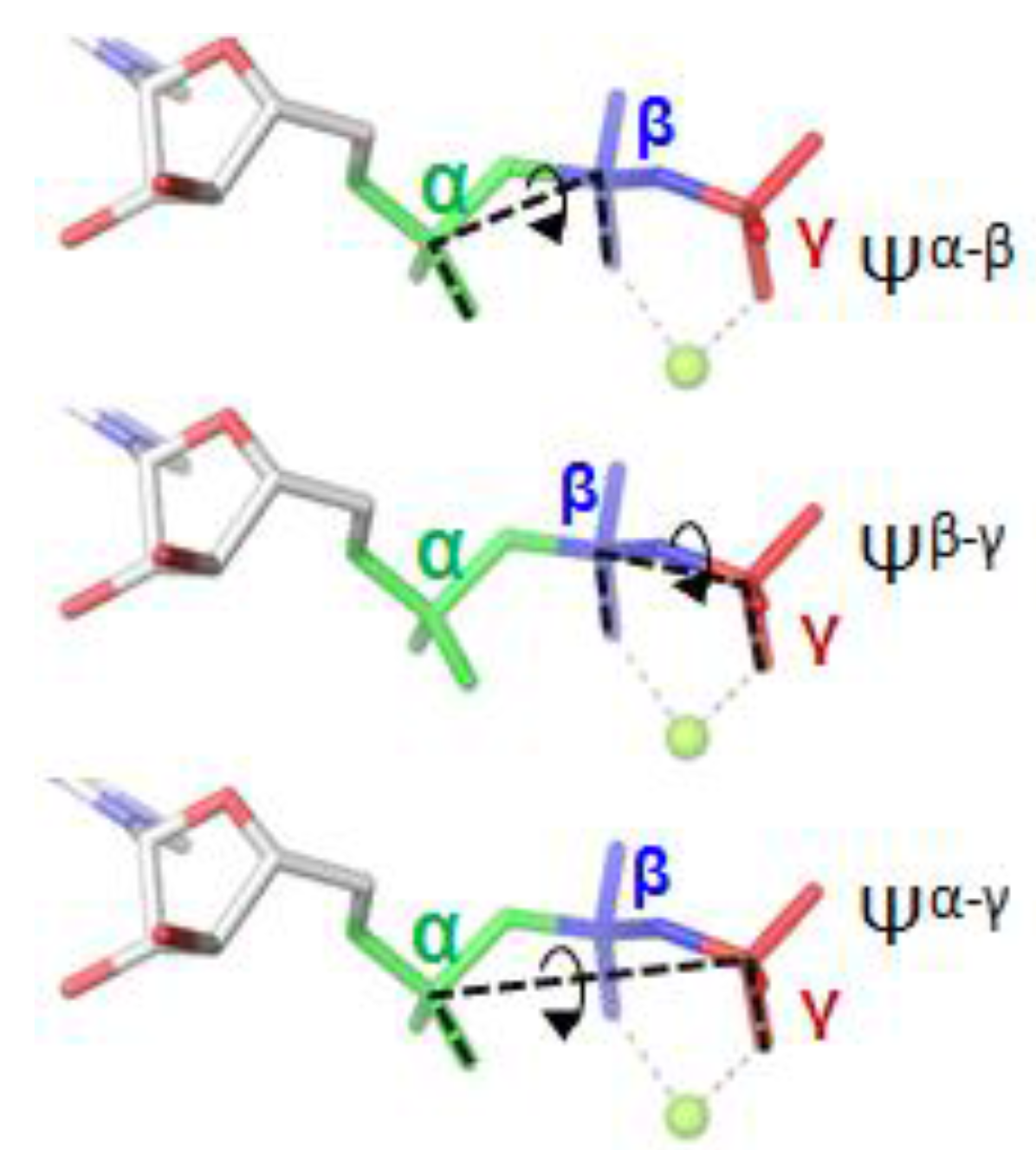
Phosphate chain conformation and dihedral angles. Dihedral angles used to describe the shape of the phosphate chain. Phosphate groups and their analogs are colored as follows: α-phosphate in green, β-phosphate in blue, and γ-phosphate in red.

**Figure S3.**
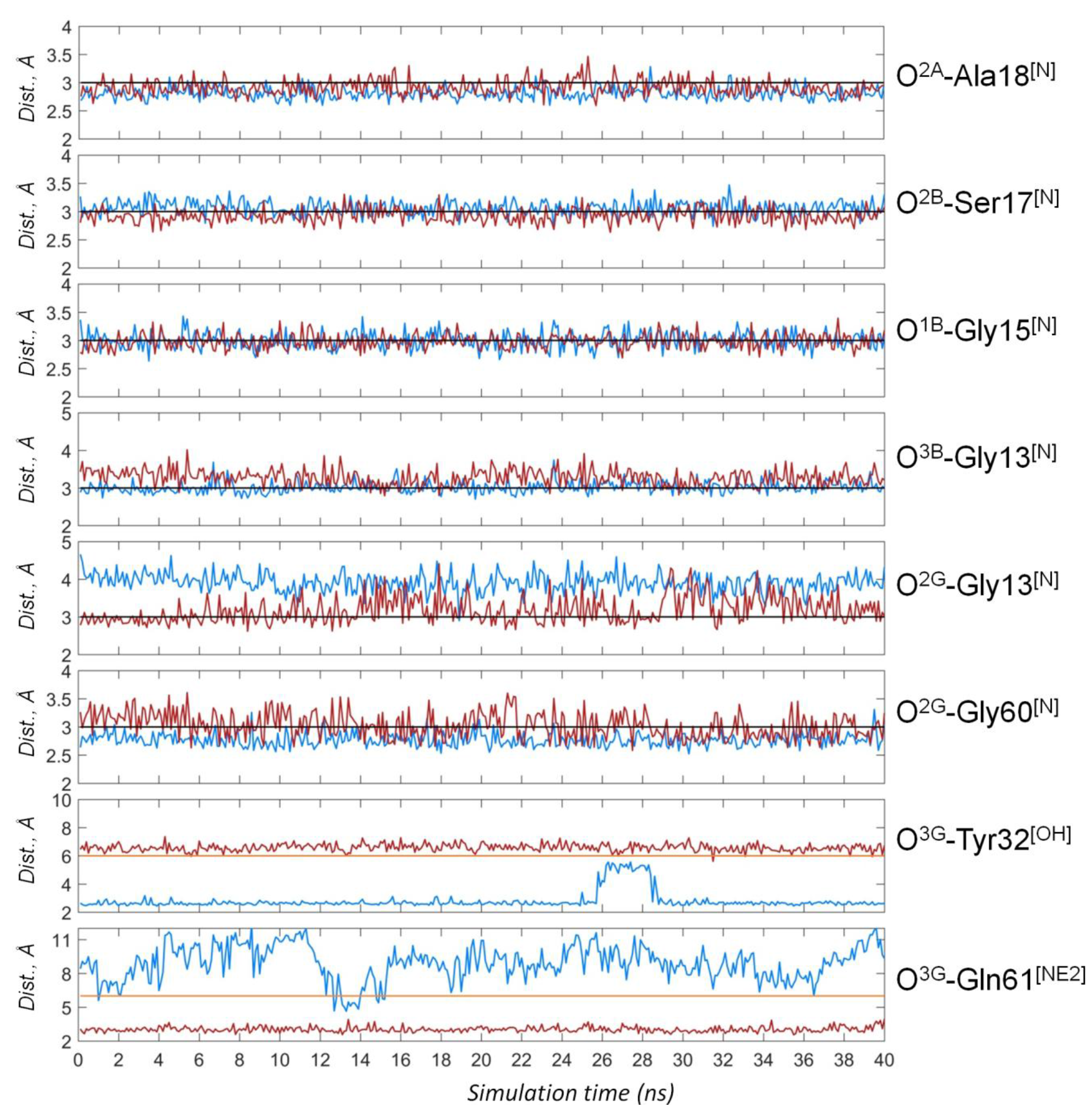
Hydrogen bonds lengths during MD simulations. Distances corresponding to the hydrogen bonds between phosphate chain oxygen atoms and surrounding amino acid residues were measured in the course of four 10-ns MD simulations. Simulation times 0-10, 10-20, 20-30 and 30-40 ns correspond to independent simulations. Blue lines show simulation of Mg-GTP-Ras, red lines are for Mg-GTP-Ras/RasGAP complex; the black lines indicate the distance of 3 Å. Distances were measured and plotted in Matlab (“MATLAB and Statistics Toolbox Release 2017a,” 2017).

**Figure S4.**
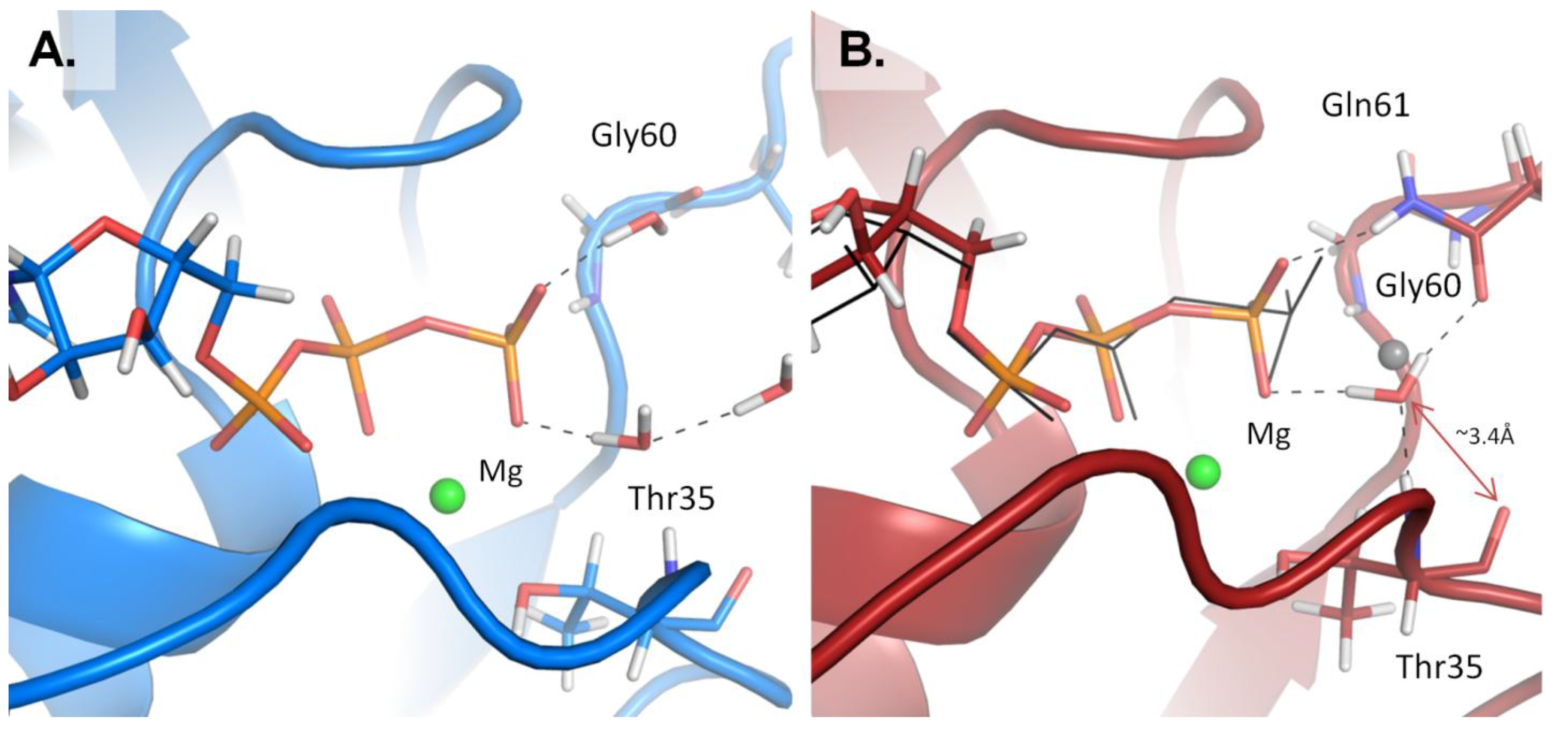
Water molecules in the active site of Ras and Ras/RasGAP. **A.** MD simulation of the Mg-GTP-Ras complex, the protein is shown as a blue cartoon, GTP and water molecules are shown as sticks, the Mg cation is shown as a green sphere, hydrogen bonds are shown as black dashed lines. **B.** The MD simulation of the Mg-GTP-Ras/RasGAP complex, the proteins are shown as a red cartoon, the GTP, water molecule and Mg^2+^ ion are colored as in panel A. The conformation of GDP-AlF_3_ complex, as taken from the crystal structure [PDB 1WQ1], is shown as black lines, the catalytic water molecule from the same structure is shown as a grey sphere.

**Figure S5.**
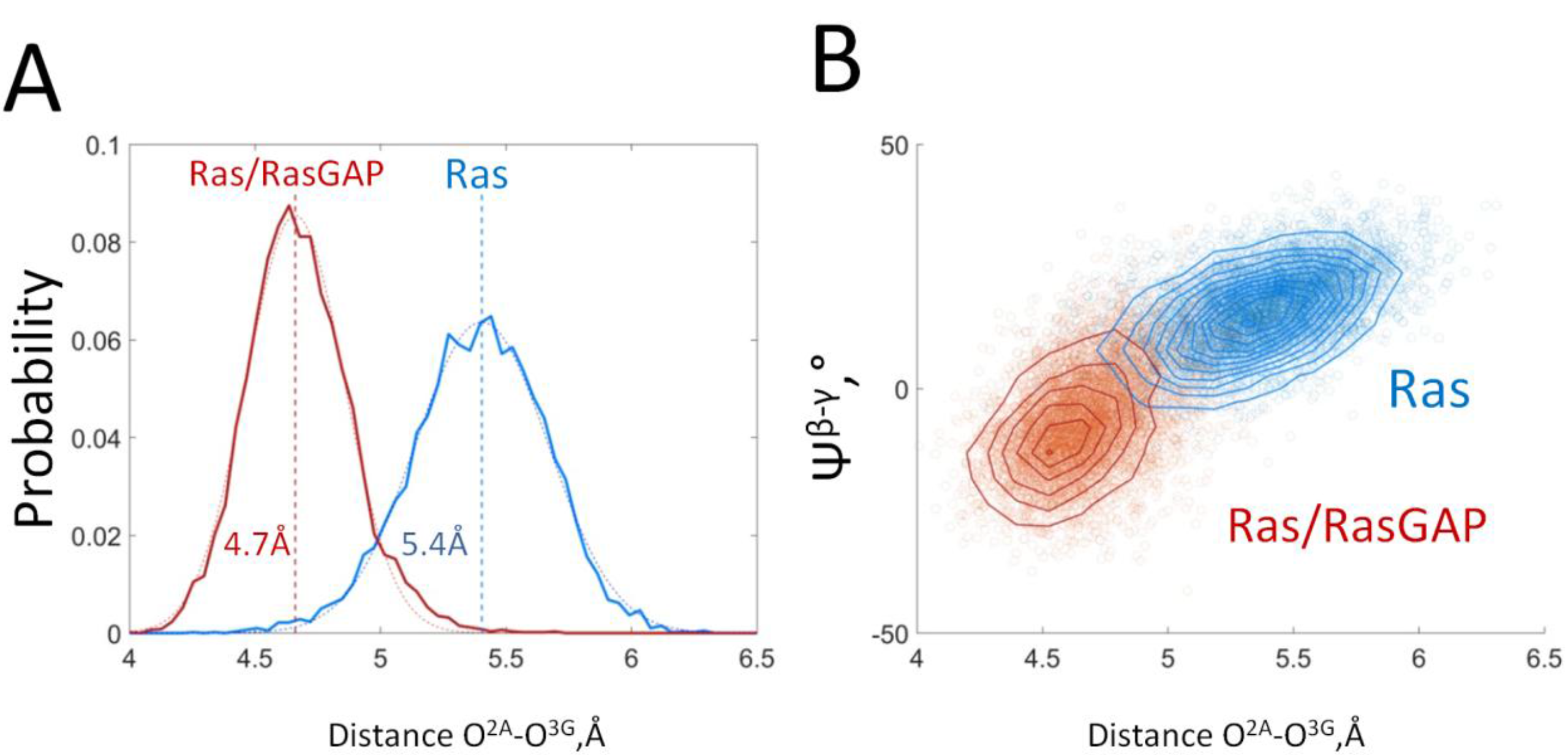
The distances between O^2A^ and O^3G^ atoms in GTP bound to Ras and Ras/RasGAP. **A.** Distribution histograms of the O^2A^-O^3G^ distance during MD simulations of Mg-GTP bound to Ras (blue) and Ras/RasGAP (red), The histograms were fitted with Gaussian in MatLab, the corresponding average values are shown as dashed lines. **B.** Correlation between the O^2A^-O^3G^ distance (X axis) and the value of dihedral angle Ψ^β-γ^ (Y axis). Individual conformations observed in MD simulations are plotted as circles; isolines are drawn to indicate points having the same distribution density.

**Figure S6.**
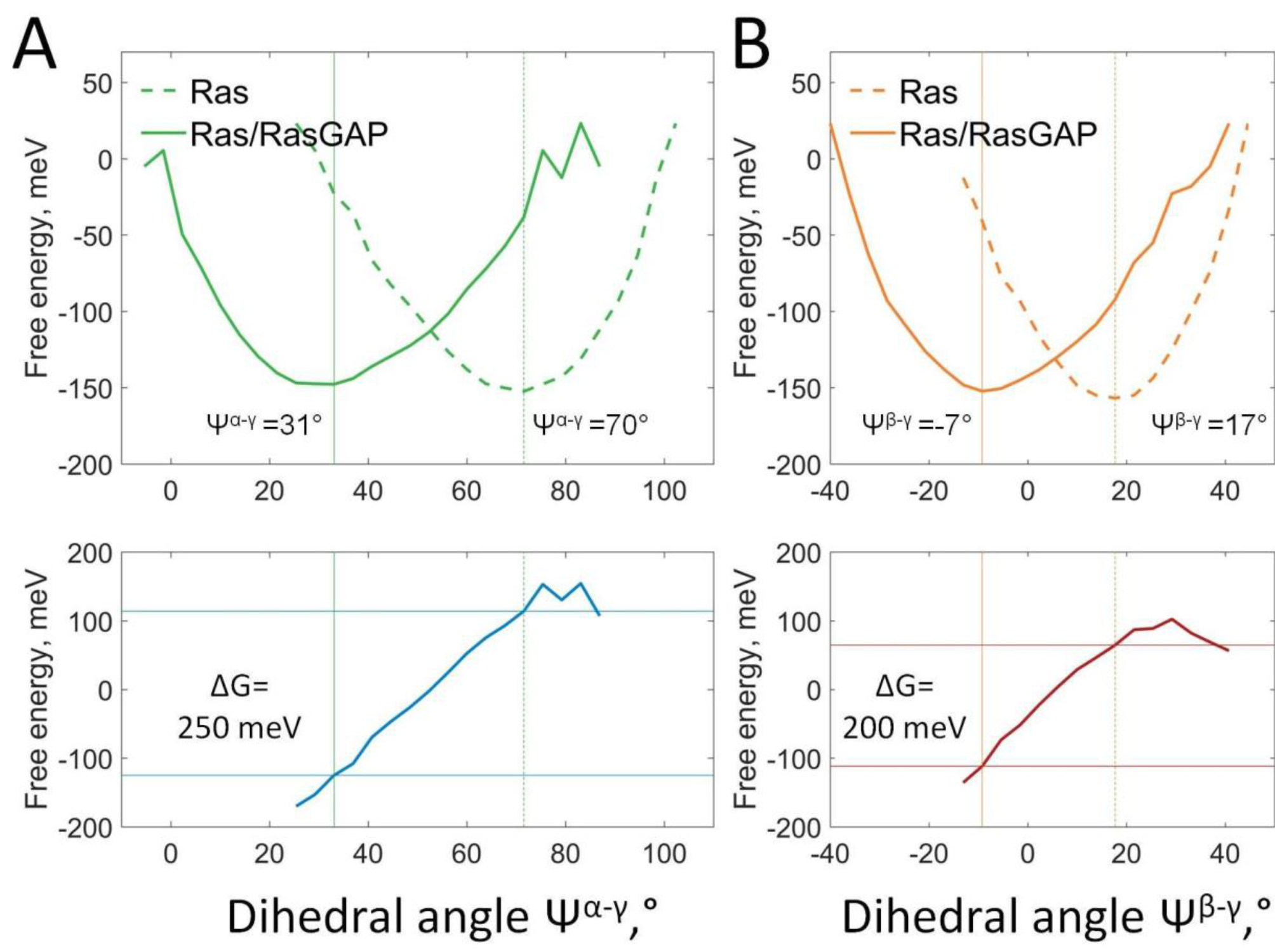
Free energy of the phosphate chain rotation in response to the binding of RasGAP to Ras as a function of dihedral angles values. Dihedral angles were measured as shown on Figure S2. The lowest energy values were set to zero. **A.** Rotation of γ-phosphate relative to α-phosphate. Top plot: Free energy calculated from probabilities (normalized frequencies) of Mg-GTP conformations plotted as function of the dihedral angle between α- and γ-phosphates. Data from four 10-ns MD simulations of Mg-GTP-Ras (green dotted line) and Mg-GTP-Ras/RasGAP (green line). Bottom plot: Free energy of rotation of γ-phosphate relative to α-phosphate, calculated as the difference between the traces in the top plot on panel A. **B.** Rotation of γ-phosphate relative to β-phosphate. Top plot: Free energy calculated from probabilities (normalized frequencies) of Mg-GTP conformations plotted as function of the dihedral angle between β- and γ-phosphates. Data from four 10-ns MD simulations of Mg-GTP-Ras (dotted orange lines) and Mg-GTP-Ras/RasGAP (orange lines) complexes. Bottom plots: Free energy of rotation of γ-phosphate relative to β-phosphate, calculated as the difference between the traces in the top plot on panel B.

### Statistical Analysis

To analyze the conformation of Mg-GTP in complex with Ras and Ras/RasGAP we measured dihedral angles between the phosphate groups. To select independent measurements among all the conformations obtained upon MD simulations, we calculated autocorrelation functions for respective angles (Fig. S7). From this plot, the correlation time was estimated at 50 ps of MD simulation time, providing 600 independent frames/measurements from the total 30 ns of simulation time for each of two systems. Only such independent frames were used upon structure analysis to calculate characteristic parameters.

**Figure S7.**
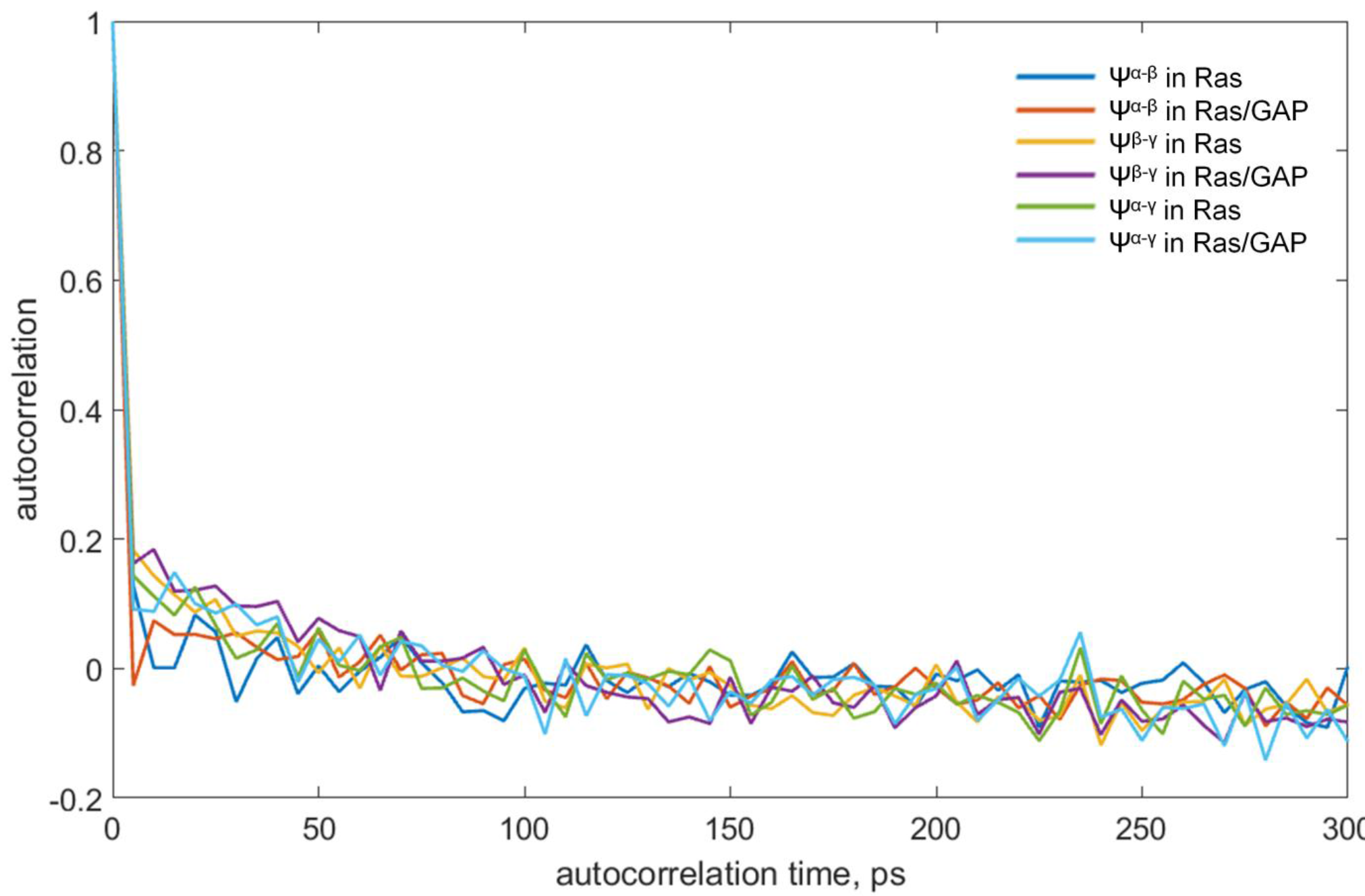
Estimation of correlation times for the measurements of dihedral angles between the phosphate groups. Autocorrelation values are plotted as a function of the autocorrelation time.

**Table S5.** Analysis of the dihedral angle measurements for the Mg-GTP in complex with Ras and Ras/RasGAP.

| Angle | Mg-GTP-Ras |  | Mg-GTP-Ras/RasGAP |  | P-value |
| --- | --- | --- | --- | --- | --- |
|  | Value | Sample size | Value | Sample size |  |
| $\Psi^{\alpha-\beta}$ | $59 \pm 6^\circ$ | 300 | $46 \pm 6^\circ$ | 300 | $10^{-97}$ |
| $\Psi^{\beta-\gamma}$ | $17 \pm 8^\circ$ | 300 | $-6 \pm 11^\circ$ | 300 | $10^{-113}$ |
| $\Psi^{\alpha-\gamma}$ | $69 \pm 10^\circ$ | 300 | $33 \pm 12^\circ$ | 300 | $10^{-170}$ |

The null hypothesis was that the dihedral angles of Mg-GTP in both complexes (Ras and Ras/RasGAP) result from normal distributions with equal mean values.
P-values were obtained from the two-sample t-tests with the null hypothesis that dihedral angles in Mg-GTP in Ras and Ras/RasGAP result from normal distributions with equal mean values.

